# N-GLYcanyzer monitoring of a microperfusion process reveals critical parameters’ impact on Trastuzumab biosimilar N-glycoform time evolution

**DOI:** 10.64898/2026.10.31.763318

**Authors:** Aditya Narvekar, Malcolm Harrison, Rahul Thakur, Padmadisha Khot, Pauline Semira, Kevin Lee, Shishir P.S. Chundawat

## Abstract

Controlling monoclonal antibody (mAb) N-glycosylation as a critical quality attribute (CQA) remains challenging in continuous and intensified biomanufacturing, where conventional end-point analyses obscure dynamic pathway responses. Here, we deployed the N-GLYcanyzer process analytical technology (PAT) for automated monitoring of a microperfusion process that reveals critical parameters’ impact on Trastuzumab biosimilar N- glycoform time evolution across a 20-day culture duration of Chinese Hamster Ovary (CHO) cells in a ∼2 mL scale-down microbioreactor perfusion platform. Using offline PAT assays combined with our integrated at-line PAT workflow, we systematically screened six perfusion process parameters: carbon source, viable cell density (VCD), perfusion rate, temperature, and pH. While nutrient limitation and thermal/acidic stress significantly disrupted N-glycans maturation, notably triggering high-mannose (Man5) accumulation and severe productivity penalties under pH 6.7 and 39 °C operation, targeted feed supplementation with galactose and MnCl consistently shifted glycoform profiles away from G0F toward galactosylated species (G1F, G1F′, G2F). Crucially, coupling this combined feed strategy with an intensified perfusion rate (1.2 VVD) achieved a sustained, desired galactosylation shift without a productivity cost. These findings demonstrate that nutrient feed composition and perfusion rate are primary levers for steering continuous intensified microperfusion processes for biologics production with desired N-glycoform states and highlight the utility of automated at-line PAT like N-GLYcanyzer for enabling real-time attribute-based control in continuous biomanufacturing.

## 1. Introduction

Monoclonal antibodies (mAbs) continue to dominate the biotherapeutic landscape, with applications spanning oncology, autoimmune disorders, and infectious disease (Walsh and Walsh, 2022). A defining feature of these molecules is the conserved N-linked glycosylation site at Asn297 in the Fc region, where the attached glycan structure directly modulates antibody effector functions, pharmacokinetics, and immunogenicity (Reusch and Tejada, 2015). The functional consequences of glycan heterogeneity at this site are well established: removal of core fucose substantially enhances binding to FcγRIIIa and amplifies antibody-dependent cellular cytotoxicity (ADCC) (Shields et al., 2002), terminal galactosylation influences complement-dependent cytotoxicity (CDC) through modulation of C1q binding (Hossler et al., 2009), and sialylation has been implicated in anti-inflammatory activity via interactions with DC-SIGN and FcγRIIB (Anthony and Ravetch, 2010). Because even modest changes in glycoform distribution can alter therapeutic potency, N-glycosylation is classified as a critical quality attribute (CQA) that must be maintained within defined limits throughout manufacturing. This requirement is especially stringent in the development of mAb biosimilars like Trastuzumab (Reusch and Tejada, 2015).

N-glycosylation of mAbs produced in Chinese hamster ovary (CHO) cells is sensitive to a range of upstream process parameters, including culture temperature, pH, dissolved oxygen (DO), osmolality, and carbon source, as well as media supplementation strategies involving galactose, manganese, and uridine (Hossler et al., 2009). However, a recent systematic review that applied a standardized glycan index framework to evaluate 78 process parameter manipulations across the published literature found that manipulations of pH, dissolved oxygen, and CO partial pressure rarely produced meaningful shifts in glycosylation, with changes of less than 5% observed for fucosylation, galactosylation, and sialylation indices (Bravo-Venegas et al., 2026). By contrast, in several cases temperature and osmolality changes notably affected galactosylation (exceeding 10%) and fucosylation (1–10%). These findings suggest that, for pH, DO, and pCO specifically, isolated process parameter adjustments are generally insufficient to meaningfully steer glycan profiles, whereas temperature and osmolality could produce notable galactosylation shifts in a subset of cases. In contrast, targeted media supplementation with uridine, manganese, and galactose has been shown to increase the galactosylation index by more than 20% across multiple CHO systems(Gramer et al., 2011), and the targeted feeding approach has demonstrated that galactose availability can be precisely controlled to tune glycosylation in continuous perfusion culture (Zhang et al., 2020). Collectively, these findings indicate that nutritional strategies, particularly carbon source manipulation and cofactor supplementation, represent the most effective levers for modulating mAb glycosylation during upstream bioprocessing.

The biopharmaceutical industry is increasingly transitioning from conventional fed-batch toward continuous and intensified perfusion processes, motivated by the potential for higher volumetric productivity, reduced facility footprint, and more consistent product quality at metabolic steady state (Coffman et al., 2021). Within this framework, miniaturized perfusion bioreactors such as the Mobius® Breez microbioreactor system (MilliporeSigma, Burlingon, MA), an automated microfluidic platform with a ∼2 mL working volume, have emerged as powerful scale-down tools that enable parallel evaluation of process conditions with substantially reduced media consumption and labor compared to bench-scale stirred-tank bioreactors (Schwarz et al., 2023). Despite these advances in perfusion platform development, the relationship between critical process parameters and glycosylation has been examined in only a small number of perfusion studies, and systematic multi-parameter screening of glycosylation dynamics in microperfusion bioreactors remains essentially unexplored. Of the 78 process parameter manipulations surveyed previously (Bravo-Venegas et al., 2026), only two were conducted under any form of perfusion operation, and none employed microperfusion bioreactors. This gap is significant because perfusion cultures maintain a distinct extracellular environment relative to batch and fed-batch processes: nutrients are delivered continuously, by-products such as lactate and ammonia are removed rather than allowed to accumulate, and a high cell density is sustained near steady state rather than following a growth-and-decline trajectory (Liang et al., 2024). Because N-glycosylation is set by cellular metabolic flux and the availability of intracellular nucleotide-sugar precursors (Burleigh et al., 2011), these differences would be expected to shape glycoform trajectories over the course of a run.

Concurrent with advances in continuous bioprocessing, process analytical technology (PAT) has become integral to upstream process development, with established tools for real-time monitoring of metabolites, cell density, and product titer (2004). Recent work has extended PAT capabilities to include automated online liquid chromatography workflows for monitoring mAb concentration, aggregation, and charge heterogeneity directly from bioreactor harvest (Dunn et al., 2023). Raman spectroscopy coupled with partial least squares chemometric modeling has also been applied to predict amino acid concentrations and, indirectly, N-glycosylation in high-density perfusion cultures (Schwarz et al., 2022), though these approaches provide correlative rather than direct glycan measurements (Li et al., 2018). For direct glycoform-resolved monitoring, (Tharmalingam et al., 2014) pioneered a real-time glycosylation monitoring framework using a micro sequential injection system coupled with UPLC, enabling automated Protein A capture, glycan release, 2-AA labeling, and HILIC separation with 12-hour temporal resolution in fed-batch culture. This concept was subsequently advanced by (Gyorgypal and Chundawat, 2022), who developed the N-GLYcanyzer, a fully automated PAT platform incorporating optimized aqueous 2-AB labeling with a nontoxic reductant (2-picoline borane), and integrated HPLC analysis within approximately 3 hours of sample draw, and validated it in fed-batch culture. Further iterations of the N-GLYcanyzer integrated instant procainamide labeling chemistry for a nearly 10-fold reduction in sample preparation time and LC-MS compatibility (Gyorgypal et al., 2023), and the platform was subsequently applied to perfusion bioreactor cultures (Gyorgypal et al., 2023). Despite these advances, no direct, glycoform-resolved PAT platform has yet been integrated with a microperfusion bioreactor to our knowledge, nor has automated glycan monitoring been combined with systematic multi-parameter process screening to evaluate glycosylation dynamics across varied culture conditions. To address this gap, our group has developed the N-GLYcanyzer, an integrated PAT platform that automates Protein A purification, N-glycan release, fluorescent labeling, and UPLC-based chromatographic separation to deliver glycoform-resolved analysis of mAb N-glycosylation within hours of sample draw (Gyorgypal and Chundawat, 2022). The platform has been demonstrated for at-line monitoring of both fed-batch and perfusion bioreactor cultures, producing glycan profiles that show strong concordance with conventional offline analytical methods (Gyorgypal et al., 2023). In the present work, we apply the N-GLYcanyzer to analyze perfusate samples from microperfusion cultures throughout the run.

In this study, we systematically investigated how multiple critical process parameters shape the time-resolved N-glycosylation profile of Trastuzumab produced in CHO microperfusion cultures over 20-day runs. The parameters evaluated included carbon source (glucose versus glucose supplemented with galactose and MnCl), viable cell density (VCD), perfusion rate (1.2 and 1.5 VVD), and temperature and pH perturbations, with daily sampling across each 20-day run to capture time-resolved glycoform trajectories. Daily offline glycan profiling using procainamide-labeled UPLC provided glycoform-resolved trajectories throughout each run, capturing dynamic shifts that cannot be resolved by conventional end-point analysis. The N-GLYcanyzer PAT workflow was additionally applied to harvest to evaluate its concordance with offline measurements and its potential for rapid glycan quality assessment of microperfusion-derived product. To our knowledge, this is the first study to combine systematic multi-parameter process screening with daily glycoform-resolved monitoring in a microperfusion bioreactor system, a culture mode that has seen growing adoption of miniaturized platforms for accelerated process development (Schwarz et al., 2023), but in which glycoform-resolved characterization has not previously been reported. Results from reported experiments are expected to clarify which parameters most strongly govern N-glycan heterogeneity for mAb produced during continuous microperfusion operations.

## 2. Materials & Methods

### 2.1 Materials

HIP-CHO basal medium was obtained from Thermo Fisher Scientific, Waltham, MA. L-methionine sulfoximine (MSX) was from Sigma-Aldrich (St. Louis, MO). D-glucose, D-galactose and manganese(II) chloride (MnCl□) were purchased from Sigma-Aldrich (St. Louis, MO). Sodium carbonate (Na CO□), used for pH control, was from VWR (Radnor, PA). Trypan blue solution (0.4%) and phosphate-buffered saline (PBS) were from Thermo Fisher Scientific (Waltham, MA). The QuadCount cell counter was from Accuris Instruments (a division of Benchmark Scientific, Sayreville, NJ). Sodium phosphate and citric acid used for the mobile phases were from VWR (Radnor, PA). PNGase F was from Bulldog Bio, Webster, NY, USA. N-lauroylsarcosine (N-LS), dithiothreitol (DTT), procainamide hydrochloride, sodium cyanoborohydride (NaBH CN), dimethyl sulfoxide (DMSO), glacial acetic acid and octanal were purchased from Thermo Fisher Scientific, Waltham, MA. Ammonium formate, formic acid, phosphate buffer components and acetonitrile were from Thermo Fisher Scientific, Waltham, MA.

### 2.2 Methods

#### 2.2.1 Cell culture and microperfusion seed preparation

A trastuzumab-expressing Chinese hamster ovary cell line engineered with the glutamine synthetase selection system (CHO-GS) was obtained from GenScript Biotech Corporation (Piscataway, NJ). Cells from a single working cell bank vial were thawed and cultured in 20 mL of CD-CHO basal medium (Thermo Fisher Scientific, Waltham, MA) with 0.1% anticlumping agent (Thermo Fisher Scientific, Waltham, MA) in a 125 mL unbaffled shake flask (VWR, Radnor, PA) at 37 °C, 5% CO□, and 130 rpm in a New Brunswick S41i CO incubator shaker (Eppendorf, Hamburg, Germany). Upon reaching a cell density of 6–8 × 10 cells/mL, cells were subcultured into a 1:1 (v/v) mixture of CD-CHO and HIP-CHO basal media (20 mL total) supplemented with 0.1% anticlumping agent and 25 μM methionine sulfoximine (MSX) to maintain GS selection pressure. Subsequent passages were performed at the same density threshold in 125 mL shake flasks to complete media adaptation, first into HIP-CHO basal medium, then into HIP-CHO basal medium containing 8 g/L glucose, with 0.1% anticlumping agent and 25 μM MSX maintained throughout. All subcultures were performed in 125 mL flasks, as the microperfusion bioreactor requires a comparatively small inoculum volume.

#### 2.2.2 Microperfusion Bioreactor Setup and Operation

Perfusion cell cultures were carried out using the Mobius® Breez microbioreactor system, an automated microfluidic platform with a nominal working volume of 2 mL. Four microbioreactors were operated in parallel per experimental run. The system comprises three interconnected 1 mL chambers separated by gas-permeable silicone membranes, of which two are filled during operation to maintain the 2 mL working volume. Mixing is achieved through pneumatic actuation of the silicone membranes, which also enables bubble-free gas exchange of O□, CO□, and air across the membrane surface. Cell retention is performed through an integrated polyethersulfone (PES) filter with a 1.2 μm pore size, which separates cell-free permeate from the culture while retaining cells within the growth chamber. The system is equipped with integrated optical sensors for dissolved oxygen (DO), pH (dual sensors for drift correction), and optical density (OD), the latter of which was used in a feedback control loop to regulate viable cell density through automated cell bleeding.

Bioreactors were inoculated at a cell density of 0.6 × 10 cells/mL for standard perfusion runs. For one intensified perfusion experiment, cells from the shake-flask culture were concentrated by centrifugation and inoculated at a seeding density of 30–40 × 10 cells/mL, with perfusion operated at 1.2 VVD. Process setpoints were maintained at 37 °C, 50% DO, and pH 7.1 ± 0.2, with pH controlled via CO in the inlet gas and supplementation of 0.5 M Na CO solution. The base perfusion medium for all runs consisted of HIP-CHO basal medium supplemented with 8 g/L glucose. For experimental conditions evaluating the effect of galactose on N-glycosylation, the perfusion medium was further supplemented with 8 g/L galactose and 1 μM MnCl□. Perfusion was operated at fixed rates of 1.2 or 1.5 reactor volumes per day (VVD), depending on the experimental condition. The target viable cell density was varied across experiments (20 x 10^6^ cells/mL or 50 x 10^6^ cells/mL) to evaluate the effects of cell density, carbon source, perfusion rate, temperature, and pH perturbation on N-glycosylation of the expressed mAb. In OD-controlled runs, cell density was held at a defined setpoint through automated bleeding triggered by the OD sensor. Sampling was performed daily by ejecting 120 μL of cell suspension from the bioreactor for offline cell count analysis. The perfusate collected in the microcentrifuge tube was used for metabolite estimation.

#### 2.2.3 Estimation of cell count, viability, and cell culture metabolites

Cell density and viability were assessed daily using a QuadCount cell counter with trypan blue exclusion. Cell suspension samples were mixed 1:1 v/v with 0.4% trypan blue solution before counting. When the viable cell density exceeded 8 × 10 cells/mL, samples were first diluted 1:5 in 1× PBS before mixing 1:1 with 0.4% trypan blue to maintain counting accuracy within the instrument’s linear range. Key cell culture metabolites, including glucose, lactate, and ammonia, were quantified from 135 μL of retained perfusate using a Nova Flex2 bioanalyzer (Nova Biomedical, Waltham, MA) equipped with the appropriate sensor card and chemistry calibration reagent.

#### 2.2.4 Protein A purification and titer estimation by UPLC

Cell-free perfusate samples were collected aseptically from each bioreactor via the sample port starting on day four for purification and titer estimation. Protein purification was performed using 0.2 mL NAb Protein A Plus Spin Columns (Thermo Scientific) loaded with 500 μL of perfusate according to the manufacturer’s protocol. Eluted fractions were analyzed by Ultra-Performance Liquid Chromatography (UPLC) using a Bio-Monolith Protein A column (Agilent Technologies) with UV detection at 280 nm, with mobile phase A being 50 mM sodium phosphate at pH 7.4 and mobile phase B being 100 mM citric acid at pH 2.8. The separation was performed at 1.0 mL/min using a gradient from 100% A from 0 to 1.5 min to 100% B from 1.5 to 3.5 min and back to 100% A to wash off the remaining elution buffer. Chromatographic peaks were manually integrated, and protein titer was estimated by comparison to a standard curve prepared from Trastuzumab reference standard samples of known concentration. A custom Python script was used to calculate protein concentration from the standard curve and to determine the volume of purified sample required for downstream glycan analysis.

#### 2.2.5 N-Glycan release and labeling using procainamide and its estimation by UPLC

Purified protein was aliquoted to obtain 100 μg for each sample, then denatured and subjected to enzymatic N-glycan release in a single-step reaction. The protein was combined with a denaturation solution containing N-lauroylsarcosine (N-LS; 10% w/v stock) and dithiothreitol (DTT; 400 mM stock) to achieve final working concentrations of 0.5% (w/v) and 20 mM, respectively. PNGase F (2 μL of 2 mg/mL) was added to catalyze N-glycan release, and the mixture was vortexed to ensure homogeneity. The sample was then incubated at 50 °C for 30 minutes to simultaneously denature the protein and facilitate rapid glycan liberation.

The labeling reagent was prepared fresh by dissolving procainamide hydrochloride (11 mg/100 µL) and sodium cyanoborohydride (NaBH CN; 6 mg/111 µL) in a solvent mixture of 70% dimethyl sulfoxide (DMSO) and 30% glacial acetic acid (AcOH). To ensure complete solubilization of NaBH CN, 30 µL of deionized water was added per 6 mg of reducing agent, and the solution was mixed thoroughly.

Following the PNGase F reaction, the procainamide labeling solution was added at a 1:1 (v/v) ratio relative to the sample. Tubes were capped and incubated at 65 °C to drive reductive amination of the reducing termini of released glycans for 3 hours.

Excess labeling reagent was removed by octanal-based liquid–liquid extraction. The labeled sample was mixed with octanal at a 1:2 (v/v) ratio and vortexed for 30 seconds. Phases were separated by centrifugation at 3,000 rpm for 1 minute (Minispin microcentrifuge, Eppendorf, Hamburg, Germany). The upper organic layer was discarded, and the resulting aqueous phase, containing the purified procainamide-labeled glycans, was collected and transferred to an HPLC vial for analysis.

Procainamide-labeled glycans were analyzed by HILIC-HPLC using an Agilent Online Monitoring 1290 HPLC system (Agilent Technologies Inc., Germany) equipped with a fluorescence detector set to an excitation wavelength of 308 nm and an emission wavelength of 359 nm. Chromatographic data acquisition and peak analysis were performed using Agilent OpenLab CDS software.

Separation was achieved on an AdvanceBio Glycan Mapping column (120 Å, 2.1 × 150 mm, 2.7 µm particle size; Agilent Technologies Inc.) using a binary gradient system. Mobile phase A consisted of 100 mM ammonium formate buffer (pH ∼4.4), and mobile phase B was acetonitrile (ACN). Ammonium formate was delivered in a linear gradient increasing from 15% to 100% over 48 minutes at a flow rate of 0.5 mL/min. Samples were maintained at 10°C in the multisampler compartment prior to injection, and the injection volume was 1 µL per run. Each labeled sample was injected in duplicate to confirm analytical reproducibility. With two independently labeled replicates per time point and duplicate injections of each, four data points were collected per condition.

#### 2.2.6 Estimation of glycans using the N-GLYcanyzer PAT workflow

The glycoform content was checked daily using the N-GLYcanyzer system via its ProSIA subunit (FIAlab Instruments, Seattle, WA), following the method described previously (Gyorgypal and Chundawat, 2022). Briefly, the perfusate from the Breez Microperfusion setup was delivered to the ProSIA system, which was integrated with a miniature, self-packed protein A column using MabSelect SuRe protein A resin (MilliporeSigma, Burlington, MA). Following mAb adsorption, the column was washed with 20 mM phosphate buffer (pH 7.2) and eluted with 0.1% formic acid. The eluate was passed through an in-line UV spectrometer (Ocean Optics, Dunedin, FL) positioned downstream of the Protein A column, measuring absorbance at 280 nm. Integrated peak area was used to calculate protein titer against a seven-point calibration curve of purified trastuzumab (MedChemExpress, Monmouth Junction, NJ). The eluted protein was then neutralized, and the glycans were labeled using the N-GLYcanyzer according to the standard procainamide labeling method (described above).

## 3. Results & Discussion

### 3.1 Detection range characterization of the N-GLYcanyzer PAT workflow for glycoform monitoring in microperfusion culture

To evaluate the detection range of the N-GLYcanyzer PAT workflow as shown in Figure 1 under microperfusion conditions, Trastuzumab samples spanning a titer range of 0.3–1.0 g/L were analyzed for individual glycoform peak areas (Table 1). All eight N-linked glycan species were successfully detected and quantified at titers of 0.5 g/L and above. The four predominant glycoforms G0F, G1F, G1F′, and G2F were consistently detected across the full 0.3–1.0 g/L titer range tested, consistent with the fucosylated glycan profile reported for CHO-produced IgG1 monoclonal antibodies, in which G0F, G1F, and G2F are the three most abundant species (Luo and Zhang, 2023). G0F yielded the highest peak area at all titers tested (3.04 ± 0.15 to 4.87 ± 0.22 LU·s), followed by G1F, G1F’, and G2F. Among the minor glycoforms, G1 fell below the detection threshold at 0.3 g/L but was reliably quantified at 0.5 g/L (0.35 ± 0.01 LU·s) and above. Titers below 0.3 g/L were not detectable under the analytical conditions employed in this study (Data not included here). Although peak areas did not scale proportionally with titer across all species, the data collectively demonstrate that the workflow provides reliable glycoform detection across a practically relevant titer window for microperfusion bioprocessing.

**Figure 1.**
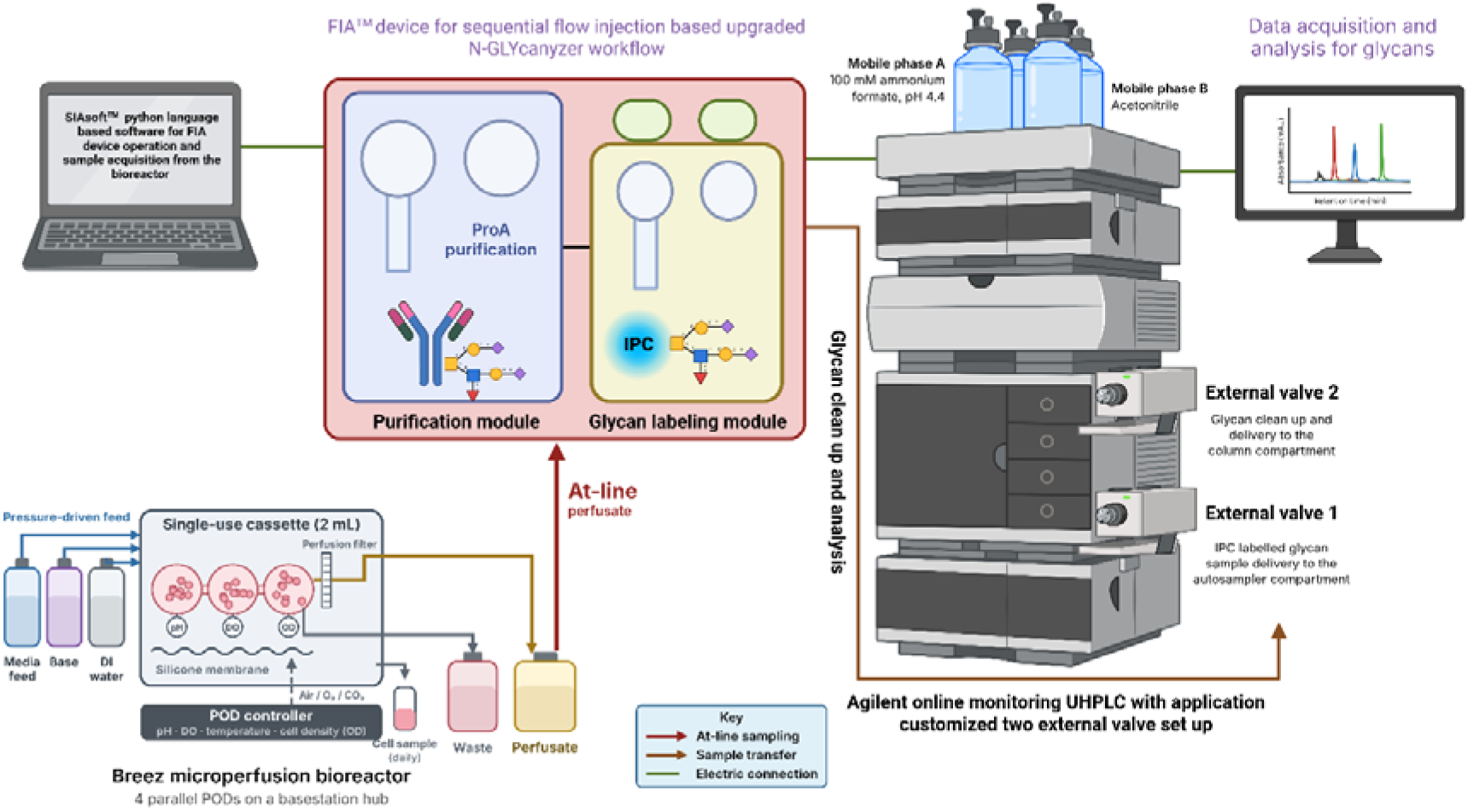
At-line N-GLYcanyzer glycan workflow integrated with the Breez microperfusion bioreactor. Cell-free perfusate is sampled at-line into the FIA device for Protein A capture, N-glycan release and IPC labeling, then analyzed on the Agilent online monitoring UHPLC with a two-external-valve set-up.

**Table 1:**
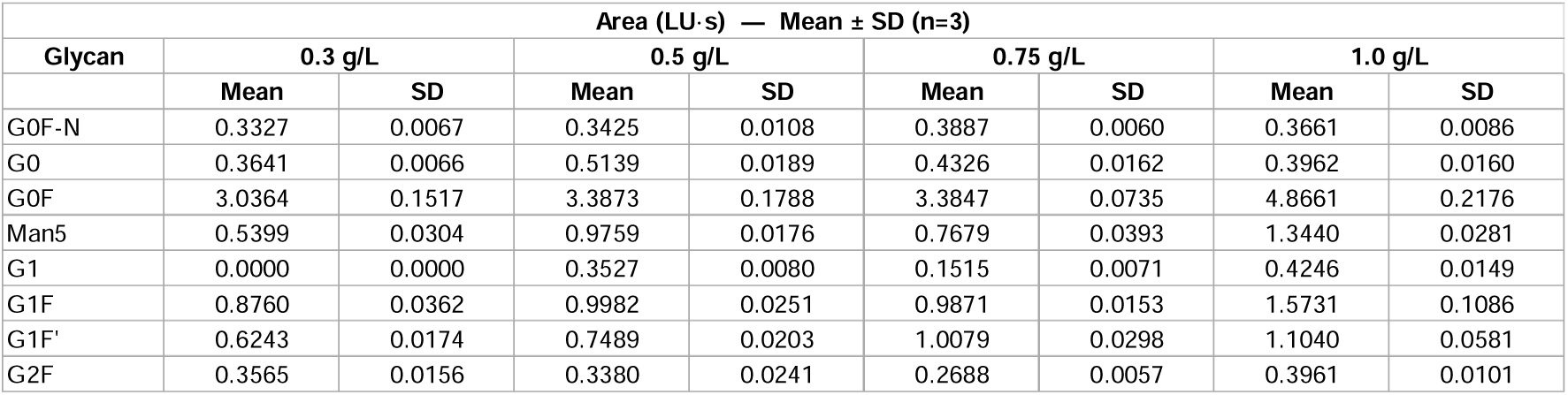
Glycan profile of trastuzumab at varying antibody titers using the N-GLYcanyzer PAT workflow. Area (LU·s) of eight N-linked glycan species (G0F-N, G0, G0F, Man5, G1, G1F, G1F’, and G2F) detected across four antibody concentrations (0.3, 0.5, 0.75, and 1.0 g/L). Data are presented as mean ± standard deviation (n = 3).

The results presented here characterize the lower titer limit of the N-GLYcanyzer PAT workflow in the microperfusion context: the four predominant glycoforms (G0F, G1F, G1F′, and G2F) remained reliably resolved down to 0.3 g/L, whereas quantification of the full eight-species panel required titers of at least 0.5 g/L. This boundary falls within the titer range observed across the microperfusion runs in this study, in which volumetric titer ranged from 0.023 to 1.30 g/L and fell at or below 0.3 g/L for extended periods in several conditions (Liang et al., 2023). The dominance of fucosylated complex-type species is consistent with the glycan microheterogeneity reported for CHO-produced trastuzumab and its biosimilars (Segu et al., 2020), in which fucosylated species constitute the large majority of the Fc glycan pool (Luo and Zhang, 2023). The non-proportional scaling of peak areas with titer, most pronounced for minor glycoforms, may reflect the narrower linear range of fluorescence detection at low analyte loads rather than a change in glycan composition. The inability to detect glycans below 0.3 g/L underscores the importance of PAT sampling timing in microperfusion workflows and indicates that N-GLYcanyzer monitoring should be initiated once the culture reaches this minimum titer threshold to ensure data quality and completeness across all glycoform species (Tharmalingam et al., 2014)

### 3.2 Impact of viable cell density on N-glycosylation at a controlled OD with glucose feed

Both conditions were maintained at a fixed perfusion rate of 1.5 VVD from Day 3 onward using glucose feed, with cell culture density independently controlled via optical density monitoring to achieve target viable cell densities of 20 × 10 and 50 × 10 cells/mL, respectively, through feedback-controlled cell bleeding (Deschênes et al., 2006). Despite matched biomass control, G0F relative abundance followed a crossover pattern between conditions as seen in Figure 2. Over Days 8–11, G0F was higher under 50 × 10 cells/mL (67.8% to 62.4%) than under 20 × 10 cells/mL (64.3% to 51.2%), a difference of up to 11%. This relationship inverted from Day 14 onward. G0F under 20 × 10 cells/mL rose to 61.6% while the 50 × 10 cells/mL condition dropped to 47.3% and then 39.8% by Day 15, the lowest value observed in either condition. By Days 19–20, the two conditions reconverged, with G0F at 49.3–49.7% (20 × 10 cells/mL) versus 50.7–53.2% (50 × 10 cells/mL). G1F remained modestly but consistently higher under 20 × 10 cells/mL across most of the culture (12.6–14.6%) compared to 50 × 10 cells/mL (9.5–13.1%). G2F showed the sharpest divergence among the major glycoforms. Both conditions tracked closely through Day 13, but 50 × 10 cells/mL spiked to 8.1% on Day 14 and 11.8% on Day 15, while 20 × 10 cells/mL remained low (2.9% and 5.7% on the same days), before both conditions returned to a comparable 3.9–6.6% range by Day 20. Among the minor glycoforms, G0F-N diverged between the two density conditions from Day 12 onward, remaining below 10% in both arms, and peaked at 9.7% under 50 × 10 cells/mL on Day 15, roughly double the 20 × 10 cells/mL trajectory, which stayed within a narrower 2.7–5.3% band. Man5 followed a similar pattern, comparable between conditions through Day 11 (∼3.8–5.7%) but diverging from Day 12 onward, reaching 9.9% under 50 × 10 cells/mL on Day 16 versus 6.2% under 20 × 10 cells/mL. G0 and G1 showed smaller, less consistent differences between conditions across the full time course.

**Figure 2.**
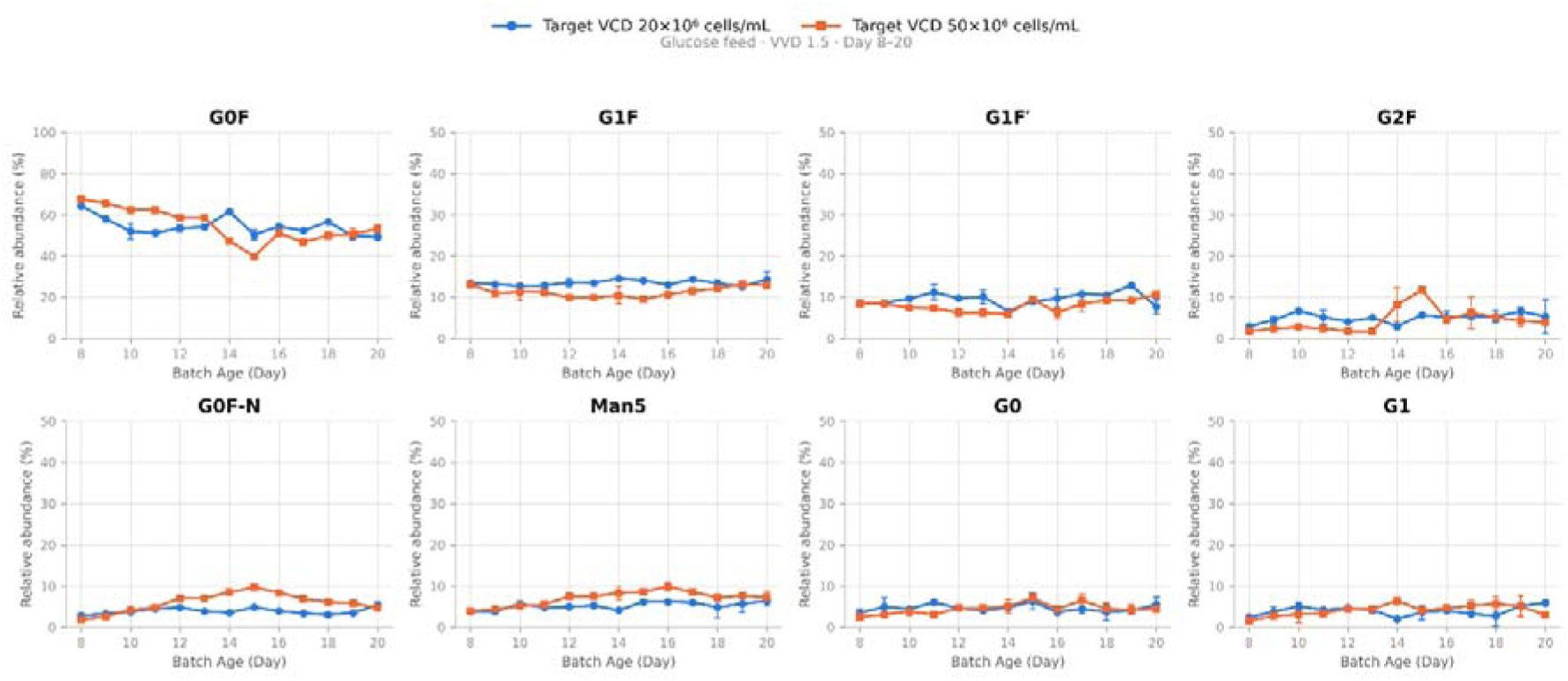
Effect of cell density on N-glycan relative abundance. Relative abundance (%) of G0F, G1F, G1F′, G2F, G0F-N, Man5, G0, and G1 over culture duration (Batch Age, Day 8– 20) under glucose feed at 1.5 VVD, comparing target viable cell density of 20×10 cells/mL (blue) versus 50×10 cells/mL (orange). G0F is plotted on a 0–100% axis; all other glycoforms are plotted on a 0–50% axis. Points represent mean ± SD.

Offline metabolite analysis showed a marked and sustained difference in glucose availability between conditions that OD control alone did not capture, as seen in Supplementary Fig. 1. Under 50 × 10 cells/mL, glucose dropped from 4.2 g/L to 0.3 g/L by Day 8 and remained glucose-limited (0.3–1.7 g/L) through Day 20 despite active glucose feeding (Mercille et al., 2000). Under 20 × 10 cells/mL, glucose never fell below 2.5 g/L, ranging 3.5–6.4 g/L across the same window; this condition was not glucose-limited at any monitored timepoint. Lactate tracked this difference closely, consistent with the strictly complementary consumption of glucose and lactate reported for CHO cultures (Ma et al., 2009). Under 50 × 10 cells/mL, lactate collapsed from 2.97 g/L to a minimum of 0.31–0.39 g/L by Days 13– 15, consistent with a glucose-starved metabolic state and with the two-phase lactate profile in which net production gives way to net consumption (Pereira et al., 2018), then reversed and rose to 1.47–2.53 g/L by Days 18–20. Under 20 × 10 cells/mL, lactate instead accumulated steadily and predictably throughout the culture, from 3.6 g/L to 4.3 g/L, the expected trend for a non-limited fed culture in which glycolytic flux remains high while residual glucose is maintained (Reinhart et al., 2015). Ammonia remained within a broadly comparable range between conditions across the culture (20 × 10 cells/mL: 0.70–1.41 mM; 50 × 10 cells/mL: 0.43–1.38 mM), with only a modest, transient elevation under 50 × 10 cells/mL at Days 13–15. Since ammonium effects on glycoform distribution are dose-dependent (Gawlitzek et al., 2000), and the two conditions did not differ systematically in ammonia exposure, that elevation postdates the onset of glycan divergence at Day 12 and followed, rather than preceded, the onset of glucose limitation on Days 7–8. Volumetric titer differed between conditions in parallel with glucose availability through Day 13 and converged thereafter. Titer under 50 × 10 cells/mL was 1.30 g/L on Day 8, roughly double the 0.63 g/L measured under 20 × 10 cells/mL on the same day, and remained elevated (0.71–0.78 g/L) through Day 13 versus 0.32–0.40 g/L under 20 × 10 cells/mL. This early productivity advantage collapsed from Day 14 onward, converging with the 20 × 10 cells/mL condition (0.41 vs 0.39 g/L on Day 14; 0.30 vs 0.28 g/L on Day 15), and both conditions plateaued in the 0.3–0.4 g/L range from Day 16 through Day 20.

The metabolite and titer data together provide run-specific context for the glycoform crossover that complements the literature-based interpretation. The 50 × 10 cells/mL condition was effectively glucose-limited from Day 7–8 onward, despite active glucose feeding and matched OD control, suggesting that cell-specific glucose demand at 50 × 10 cells/mL outpaced the delivery rate calibrated for a fixed biomass setpoint (Stepper et al., 2020). This sustained nutrient limitation coincided with an initial titer advantage under 50 × 10 cells/mL through Day 13, consistent with higher volumetric productivity from greater biomass even under per-cell glucose stress, though glucose-limited steady states have also been reported to raise cell-specific productivity (Becker et al., 2019) and the two contributions cannot be separated from volumetric titer alone. This advantage eroded sharply from Day 14 onward, converging with the 20 × 10 cells/mL condition by Day 14–17, closely aligned with the onset of the G0F crossover, the G2F spike, and the sustained rise in Man5 and G0F-N under 50 × 10 cells/mL. This timing is consistent with sustained glucose limitation from Day 7–8, rather than an acute Day-14 trigger, contributing to both the erosion of the early titer advantage under 50 × 10 cells/mL and, from Day 12 onward, the shift in glycoform distribution indicated by rising Man5 and G0F-N alongside the G0F/G2F crossover, consistent with the lag between the onset of nutrient limitation and its glycan consequences in extended CHO cultures (Wong et al., 2004).

Glucose limitation impairing nucleotide sugar donor availability and glycosyltransferase activity in the Golgi is the better-supported mechanism here, consistent with the general sensitivity of N-glycosylation to glucose and nucleotide-sugar precursor supply reported in the literature (Ivarsson et al., 2014), with quantitative glucose-depletion studies in CHO reporting reduced intracellular nucleotide and nucleotide-sugar pools alongside a galactosylation index below 0.43 and up to 51% non-glycosylated product (Villacrés et al., 2015), and with reports that reduced glycolytic flux lowers intracellular UDP-GlcNAc in parallel with decreased sialylation and antennarity (Burleigh et al., 2011). The direction of this effect is not universal, however, as glucose starvation in fed-batch culture has been reported to increase galactosylated glycoforms in some systems (Fan et al., 2015); the present dataset therefore serves as the primary evidence for this interpretation. Glycosylation flux analyses of perfusion CHO cultures similarly show that dynamic shifts in galactosylation activity were driven primarily by changes tied to cell-specific productivity and metabolic state (Hutter et al., 2018). Ammonia accumulation, previously implicated in glycan processing defects at high viable cell density in CHO perfusion culture (Karst et al., 2017), does not appear to be a major driver in this dataset: absolute ammonia concentrations remained modest and broadly comparable between conditions throughout, with only a small, transient elevation under 50 × 10 cells/mL at Days 13–15 that likely contributed marginally at most alongside the much larger glucose effect.

The reconvergence of G0F between conditions by Days 19–20 coincides with a partial metabolic recovery signal under 50 × 10 cells/mL. Lactate turns from net consumption back to net production over Days 18–20 (Hartley et al., 2018), even though glucose itself never returns to a non-limiting level. Titer over this same window plateaus in the 0.3–0.4 g/L range for both conditions, indicating that productivity stabilized rather than continuing to decline under 50 × 10 cells/mL. Taken together, this points to a coherent narrative across the culture: an initial productivity advantage under 50 × 10 cells/mL driven by higher biomass, a metabolic and glycosylation crisis from Days 12–16 driven predominantly by sustained glucose limitation with a modest ammonia contribution, and a late partial recovery, evident in both the lactate and titer trends, that restores a comparable glycoform distribution without fully resolving the underlying glucose limitation. This has a direct process-engineering implication: at a fixed OD setpoint, the glucose feed strategy calibrated for 20 × 10 cells/mL was evidently insufficient for 50 × 10 cells/mL, and matching cell-specific (rather than volumetric) glucose/media delivery would likely be necessary to avoid this transient nutrient-limitation window in future intensified perfusion runs at higher target densities, consistent with reports that cell-specific perfusion rate governs cellular metabolism and mAb N-glycosylation in high-density perfusion culture (Schwarz et al., 2022).

### 3.3 Impact of feed on Trastuzumab N-glycosylation at controlled cell density

Both conditions were maintained at a fixed perfusion rate of 1.5 VVD from Day 3 onward, with culture density independently controlled via online optical density monitoring to achieve a target viable cell density of 50 × 10 cells/mL with glucose and glucose+galactose+MnCl feed (Deschênes et al., 2006). G0F declined progressively under glucose feed across the culture, from 67.8% to a minimum of 39.8%, before partially recovering to 50.0–53.2% over Days 16–20. Under glucose+galactose+MnCl feed, G0F was substantially lower from the outset, at 41.5% on Day 8, and remained in a 27–38% range through most of the culture, aside from a single elevated value of 59.2% on Day 14. G1F was markedly and consistently higher under glucose+galactose+MnCl feed (13.5–26.6% across Days 8–20) than under glucose feed (9.9–13.1%), roughly double for most of the window. G1F’ showed the same pattern: 12.4–17.0% under glucose+galactose+MnCl feed versus 5.9–10.7% under glucose feed. G2F was also substantially elevated under glucose+galactose+MnCl feed (6.9–13.3%) relative to glucose feed (1.7–8.1%, with the higher end concentrated in the second half of the culture). Man5 followed a broadly similar trajectory under both conditions (glucose feed: 3.8–9.9%; glucose+galactose+MnCl feed: 3.7–11.7%), rising over the culture in both cases with only a modest difference in ceiling between conditions.

**Figure 3.**
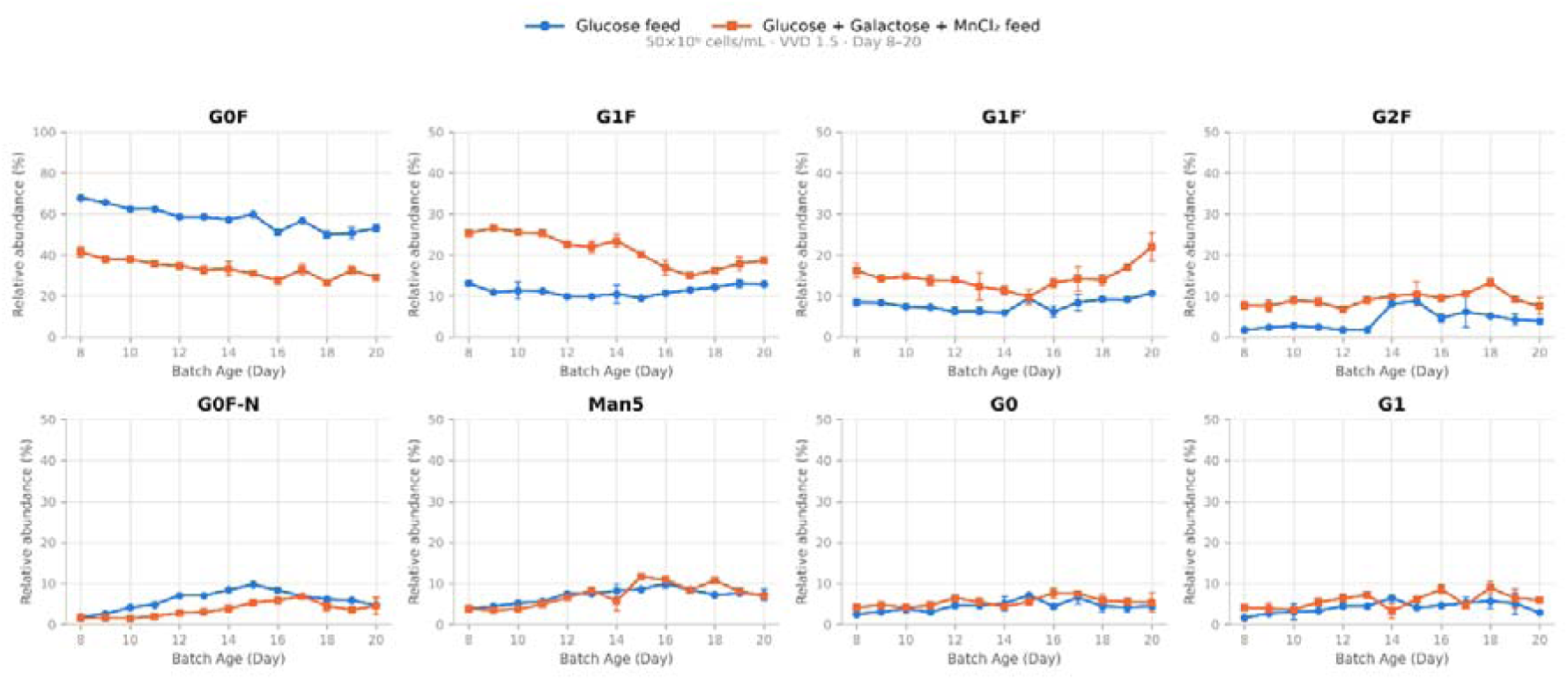
Effect of feed composition on N-glycan relative abundance. Relative abundance (%) of G0F, G1F, G1F′, G2F, G0F-N, Man5, G0, and G1 over culture duration (Batch Age, Day 8–20) at 50×10 cells/mL and 1.5 VVD, comparing glucose feed (blue) versus glucose + galactose + MnCl feed (orange). G0F is plotted on a 0–100% axis; all other glycoforms are plotted on a 0–50% axis. Points represent mean ± SD.

Glucose availability was low under both conditions across the monitored window. Residual glucose concentration for glucose feed alone ranged between 0.31–1.71 g/L, while for glucose+galactose+MnCl feed it was lower still, at 0.16–0.40 g/L throughout, indicating a more severe glucose limitation under the combined feed at this cell density and perfusion rate, consistent with the limitation of productivity by glucose availability in high-density perfusion culture (Mercille et al., 2000). Lactate followed a comparable pattern, remaining low under both conditions but generally lower under glucose+galactose+MnCl feed (0.03– 1.83 g/L) than under glucose feed (0.31–2.97 g/L), consistent with the strictly complementary consumption of glucose and lactate in CHO culture (Ma et al., 2009). Ammonia diverged substantially between conditions. Under glucose feed, ammonia remained modest throughout (0.43–1.38 mM). Under glucose+galactose+MnCl feed, ammonia rose steadily over the culture, from approximately 1.9 mM to a peak of 4.85 mM, remaining elevated (3.1–4.2 mM) through Day 20, three to four times the level under glucose feed for much of the culture.

Volumetric titer was consistently higher under glucose feed than under glucose+galactose+MnCl feed across nearly the entire monitored window. Glucose feed titer ranged 0.71–1.30 g/L over Days 8–13, compared to 0.31–0.73 g/L under glucose+galactose+MnCl feed over the same days. This gap narrowed but did not close over the remainder of the culture: 0.32–0.62 g/L (glucose feed) versus 0.18–0.44 g/L (glucose+galactose+MnCl feed) across Days 17–20 (Supplementary Fig. 2).

The substantial and sustained elevation of G1F, G1F’, and G2F under glucose+galactose+MnCl feed is consistent with the established mechanism by which galactose and manganese supplementation increase intracellular UDP-galactose availability and manganese-dependent galactosyltransferase activity, driving flux through the galactosylation pathway and shifting the glycan profile away from G0F (Gramer et al., 2011), a mechanism reviewed in the context of nucleotide-sugar precursor supplementation of growth medium (Blondeel and Aucoin, 2018) and directly observable in time-course monitoring of manganese-supplemented CHO cultures, in which galactosylation was higher than in controls while fucosylation and mannosylation remained unchanged (Tharmalingam et al., 2014). The magnitude of this shift here, with G1F approaching or exceeding double the glucose-feed value for most of the culture, indicates a strong and consistent substrate-driven effect on galactosylation at this cell density and perfusion rate. The marked elevation of ammonia under glucose+galactose+MnCl feed, reaching three to four times the glucose-feed concentration for much of the culture, is a notable feature of this comparison. Elevated ammonia has been shown to inhibit glycan processing at high viable cell density in CHO perfusion culture (Karst et al., 2017), raising the possibility that ammonia accumulation contributes to the glycoform shift observed here alongside the direct substrate-driven galactosylation effect. Ammonium concentrations in the low millimolar range, comparable to those reached here, have been reported to reduce terminal galactosylation and sialylation in a dose-dependent manner through a pH-mediated reduction in glycosyltransferase activity (Gawlitzek et al., 2000), and ammonia has been proposed to act by neutralizing Golgi pH and shifting glycosyltransferases toward less active forms (Blondeel and Aucoin, 2018), with the integrated ammonium level over a culture linked to the G0F/G1F/G2F distribution via intracellular pH regulation of β-1,4-galactosyltransferase (McCracken et al., 2014). Distinguishing the relative contribution of each mechanism would require additional data, for example testing whether the magnitude of the G1F/G2F shift tracks more closely with ammonia concentration or with galactose/MnCl dosing directly, but the concurrent rise in both ammonia and Man5 under glucose+galactose+MnCl feed is at least consistent with ammonia playing a role beyond substrate availability alone (Fan et al., 2014). The lower titer sustained under glucose+galactose+MnCl feed throughout the culture, alongside its more severe glucose limitation and markedly elevated ammonia, indicates that the galactosylation shift achieved under this feed strategy came at a consistent productivity cost at this cell density and perfusion rate, consistent with the reported impact of byproduct accumulation on product quality and protein production (Liang et al., 2024). This is a relevant consideration for process design: achieving the desired shift toward higher-order galactosylated species via glucose+galactose+MnCl supplementation was not productivity-neutral under these 14 conditions, and any implementation of this feed strategy at comparable density and perfusion rate should account for the accompanying reduction in titer.

### 3.4 Impact of intensified perfusion process on Trastuzumab N-glycosylation

Unlike the transient crossover pattern observed in the viable cell density comparison, the glucose-feed and glucose+galactose+MnCl -feed conditions at 1.2 VVD showed a large, sustained separation in G0F relative abundance across the entire Day 8–20 window, with no inversion. G0F under glucose feed (n = 2 bioreactors) remained in the 48–60% range throughout, fluctuating between 53.5% and 59.8% for most of the culture before declining to 48.3% by Day 20. Under glucose+galactose+MnCl feed (n = 2 bioreactors), G0F was consistently and substantially lower, ranging from 34–43% across the same window, a gap of 15–20 percentage points from the glucose-feed condition at nearly every timepoint.

**Figure 4.**
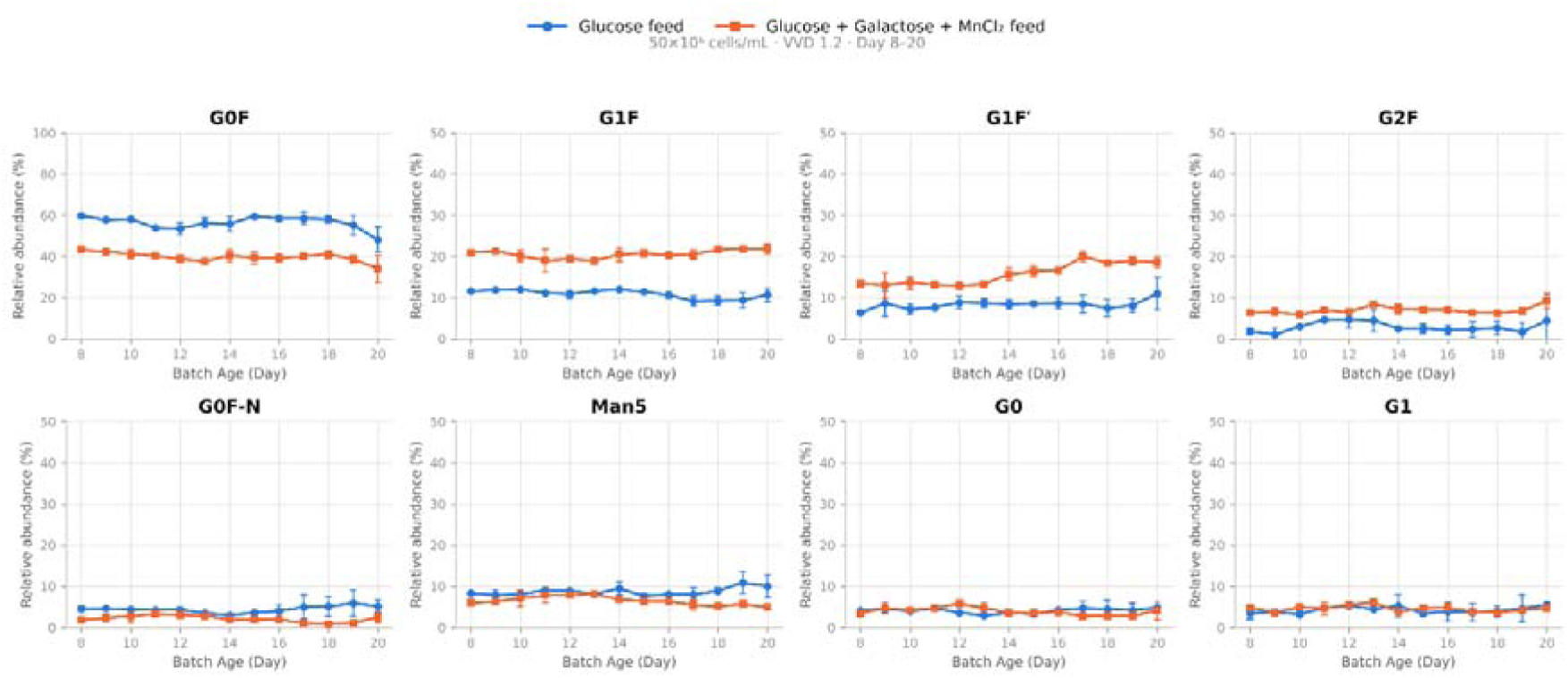
Effect of feed composition on N-glycan relative abundance under intensified perfusion. Relative abundance (%) of G0F, G1F, G1F′, G2F, G0F-N, Man5, G0, and G1 over culture duration (Batch Age, Day 8–20) at 50×10 cells/mL and 1.2 VVD, comparing glucose feed (blue) versus glucose + galactose + MnCl feed (orange). G0F is plotted on a 0–100% axis; all other glycoforms are plotted on a 0–50% axis. Points represent mean ± SD of biological duplicates.

This depletion in G0F under glucose+galactose+MnCl feed was matched by a large, sustained increase in G1F, which ranged from 19.0% to 21.9% across Days 8–20 compared to 9.2–12.0% under glucose feed, roughly double throughout. G1F’ showed an even larger relative effect and a clear upward trend specific to the glucose+galactose+MnCl -fed condition: it rose from 13.4% to a peak of 20.0% and remained elevated (18.5–18.9%) through Day 20, while under glucose feed it stayed within a comparatively narrow 6.4–11.0% band with no clear directional trend. G2F was also modestly but consistently elevated under glucose+galactose+MnCl feed (5.9–9.2%) relative to glucose feed (1.9–4.7% for most of the culture, aside from a single Day 20 spike to 8.9%). Man5 showed the opposite pattern from the galactosylated species: it was higher under glucose feed (7.6–10.9%, rising toward the end of culture) than under glucose+galactose+MnCl feed (4.9–8.1%, trending downward after Day 13). G0F-N and G0 were broadly comparable in magnitude between conditions, with G0F-N somewhat higher under glucose feed in the later culture period (up to 5.9% by Day 19 vs. 0.9–2.5% under glucose+galactose+MnCl feed from Day 17 onward).

Glucose availability differed between feed conditions in both magnitude and timing. Under glucose feed, glucose concentration declined from 3.46 g/L to a glucose-limited range (0.18– 0.22 g/L) from Day 15 through Day 19, with a partial rise to 0.55 g/L by Day 20. Under glucose+galactose+MnCl feed, glucose concentration started substantially higher (5.18– 7.10 g/L, Days 8–10) and remained non-limiting through Day 13 (>2.5 g/L), declining to a comparably glucose-limited range (0.30–0.44 g/L) only from Day 17 through Day 19, two days later than the glucose-feed condition. Lactate under glucose+galactose+MnCl feed was generally higher through the mid-to-late culture period (4.3–5.7 g/L, Days 13–19) than under glucose feed (1.6–3.9 g/L over the same window), consistent with greater total carbon flux from the combined feed. Ammonia remained within a broadly similar low range for both conditions (glucose feed: 0.62–2.34 mM; glucose+galactose+MnCl feed: 0.56–1.38 mM), with a single elevated point under glucose feed at Day 16 (2.34 mM) not mirrored in the glucose+galactose+MnCl -feed condition. Volumetric titer was comparable between conditions through Day 13 (glucose feed: 0.19–0.35 g/L; glucose+galactose+MnCl feed: 0.23–0.29 g/L). Glucose feed titer rose to a transient peak at Days 15–16 (0.55 and 0.65 g/L, respectively), exceeding glucose+galactose+MnCl feed over the same days (0.37 and 0.59 g/L). From Day 17 onward, glucose+galactose+MnCl feed titer exceeded glucose feed and remained higher through the end of the monitored window: 0.655 vs. 0.524 g/L, 0.955 vs. 0.686 g/L, 0.633 vs. 0.508 g/L, and 0.403 vs. 0.254 g/L. Replicate variability within each condition was substantial at several timepoints, most notably Days 15–18, where individual bioreactor titers diverged by up to twofold from their duplicate partner (Supplementary Fig. 3).

The magnitude and consistency of the G0F-to-G1F/G1F’/G2F shift under glucose+galactose+MnCl feed, sustained across the full monitoring window without the transient crossover seen in the density comparison, are consistent with a direct substrate-driven effect on the galactosylation pathway rather than a stress-associated or nutrient-limitation-driven mechanism, although glucose limitation developed late in both conditions (from Day 15 and Day 17, respectively) and cannot be excluded as a contributor. This aligns with the established mechanism by which galactose and manganese supplementation increase intracellular UDP-galactose availability and manganese-dependent galactosyltransferase activity, respectively, driving flux through the galactosylation pathway and increasing antibody galactosylation via a shift from G0F to G1F (Gramer et al., 2011), a mechanism reviewed in the context of nucleotide-sugar precursor supplementation of growth medium (Blondeel and Aucoin, 2018) and directly observable in time-course monitoring of manganese-supplemented CHO cultures, in which galactosylation was higher than in controls while fucosylation and mannosylation remained unchanged (Tharmalingam et al., 2014). Because the present feed strategy included both galactose and MnCl —the two supplements shown to act synergistically with uridine—alongside the base glucose feed, the present dataset is a closer match to that mechanism than a galactose-only supplement would be, though it omits the uridine component of the original UMG system. The sustained rise in G1F’ and the modest but consistent elevation in G2F beyond the primarily G0F-to-G1F shift reported in the original UMG studies may reflect this compositional difference, or differences between continuous perfusion and fed-batch culture. The lower Man5 under glucose+galactose+MnCl feed, trending downward over the culture, contrasts with the density-driven Man5 elevation observed in the earlier comparison: here, higher galactosylation coincides with lower, not higher, high-mannose content, consistent with efficient forward processing through the Golgi under a substrate-replete, non-nutrient-limited galactosylation pathway, in contrast to the incomplete-processing signature associated with metabolic stress in the density comparison.

The delayed onset of glucose limitation under glucose+galactose+MnCl feed, three to four days later than under glucose-only feed, reflects the higher total sugar supplied by the combined feed. Titer trends indicate that the sustained galactosylation shift under glucose+galactose+MnCl feed did not come at a productivity cost: titer was comparable to that under glucose feed through Day 16 and exceeded that under glucose feed from Day 17 through Day 20. This contrasts with reports that substituting glucose with galactose alone can adversely affect productivity and titer in CHO culture (Karst et al., 2017). At 1.2 VVD, supplementing the glucose feed with galactose and MnCl□, rather than substituting glucose outright, produced both a more extensive galactosylation shift, with G1F, G1F’, and G2F all elevated rather than G1F alone, and a titer advantage in the second half of the culture, suggesting that the inclusion of MnCl alongside galactose, consistent with the previously reported synergistic effect (Gramer et al., 2011), and supplementation rather than substitution, may together avoid the productivity trade-off previously reported for galactose-based glycoengineering strategies. This interpretation should be weighed against the substantial replicate-to-replicate variability observed within each feed condition at Days 15– 18, which warrants a larger sample size or tighter process control to confirm before the titer advantage is presented as a generalizable conclusion.

### 3.5 Impact of temperature perturbation on Trastuzumab N-Glycosylation

Both feed conditions were subjected to a temperature upshift from 37 °C to 39 °C beginning on Day 8. Under glucose feed, G0F declined progressively across the culture, from 60.1– 63.5% (Days 8–11) to 39.6–43.5% (Days 18–20), a gradual decline concentrated in the second half of the culture. Under glucose+galactose+MnCl feed, G0F declined earlier and more steeply: from a comparable 58.3–62.0% (Days 8–10), it fell to 45.9% by Day 11 and 39.7% by Day 12, roughly a week earlier than the equivalent decline under glucose feed, continuing to decline to 28.6–30.6% during Days 17–20.

G1F remained comparatively stable under glucose feed (12.98–15.44% across the full window). Under glucose+galactose+MnCl feed, G1F rose sharply from 15.4% to a peak of 25.3% and remained elevated (18.4–23.5%) through Day 20. G1F’ followed a similar but more modest pattern: stable at 8.4–10.3% under glucose feed, rising from 10.4% to 13.9–14.4% by Day 11 and remaining at that elevated level under glucose+galactose+MnCl feed through Day 20. G2F rose over the course of the culture under both conditions, consistent with a temperature-associated trend independent of feed: from 2.5–3.1% (Days 8–9) to 7.1– 9.9% (Days 18–20) under glucose feed, and from 2.7–3.3% (Days 8–9) to 9.3–12.2% (Days 18–20) under glucose+galactose+MnCl feed, a modestly higher ceiling under the combined feed. Man5 also rose over the course of the culture under both conditions and to a broadly similar extent (glucose feed: 4.7–5.3% early, 9.3–10.6% late; glucose+galactose+MnCl feed: 5.3–5.5% early, 5.5–9.2% late), indicating this trend was primarily temperature-associated rather than feed-dependent, although increases in Man5 with extended culture duration have also been reported as a general phenomenon in CHO cultures, independent of temperature (Fan et al., 2015).

**Figure 5.**
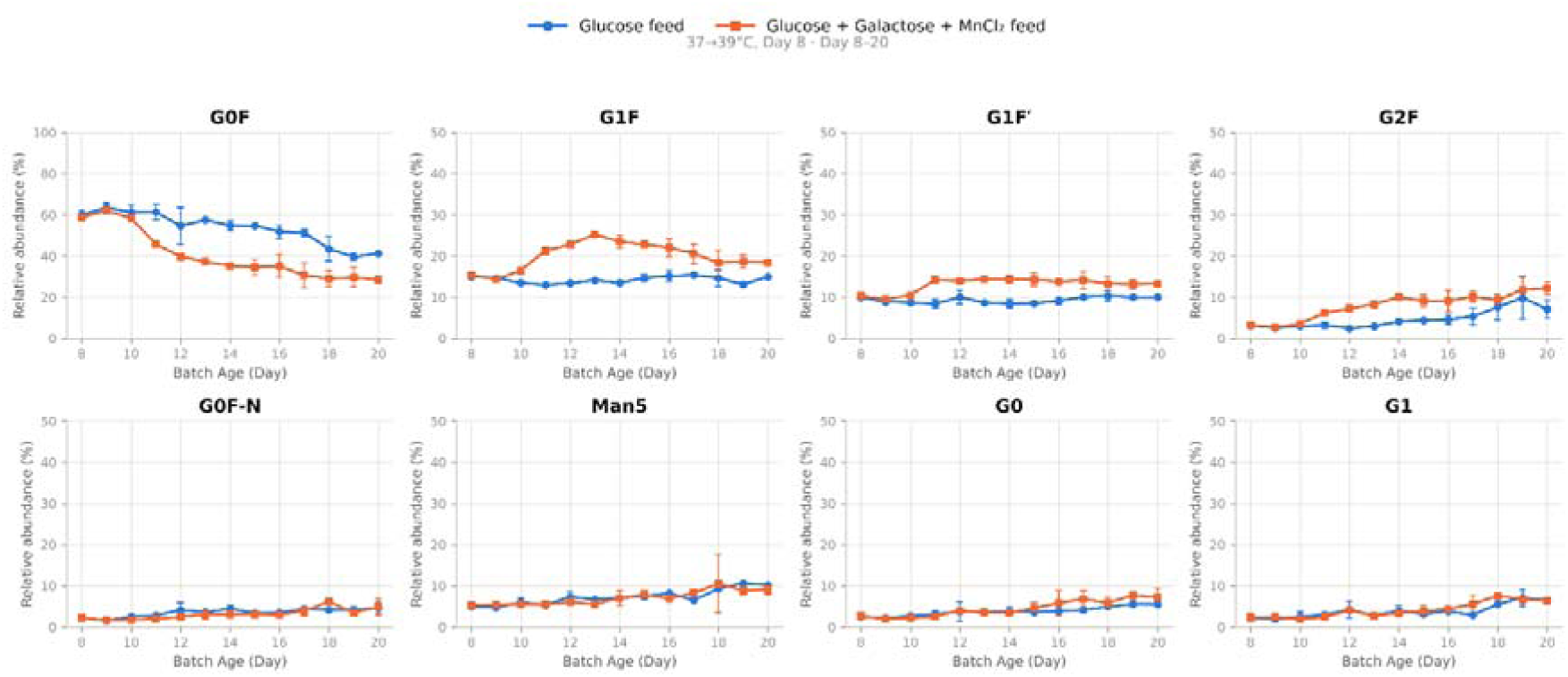
Effect of feed composition on N-glycan relative abundance under temperature upshift. Relative abundance (%) of G0F, G1F, G1F′, G2F, G0F-N, Man5, G0, and G1 over culture duration (Batch Age, Day 8–20) following a temperature shift from 37°C to 39°C on Day 8, comparing glucose feed (blue) versus glucose + galactose + MnCl feed (orange). G0F is plotted on a 0–100% axis; all other glycoforms are plotted on a 0–50% axis. Points represent mean ± SD of biological duplicates.

Glucose feed maintained higher glucose availability than glucose+galactose+MnCl feed throughout the monitored window (2.0–4.7 g/L vs. 1.2–3.0 g/L, respectively), the reverse of the pattern observed in the corresponding comparison at standard temperature, where the combined feed provided more total sugar and delayed glucose limitation. Lactate was broadly comparable between conditions (3.3–5.6 g/L under glucose feed; 3.1–6.6 g/L under glucose+galactose+MnCl feed), with no consistent separation. Ammonia was modestly and fairly consistently higher under glucose feed (1.2–3.9 mM) than under glucose+galactose+MnCl feed (0.5–2.0 mM) across most of the culture. Volumetric titer declined steadily under both conditions across the monitored window, consistent with a temperature-stress-associated loss of productivity. Glucose feed titer declined from 0.485 g/L to 0.043 g/L. Glucose+galactose+MnCl feed titer was lower than glucose feed at every timepoint, declining from 0.228 g/L to 0.041 g/L, converging with glucose feed only at the final timepoint (Supplementary Fig. 4).

The progressive decline in G0F and corresponding rise in G1F, G1F’, G2F, and Man5 observed under glucose feed over the course of the temperature-upshifted culture is consistent with reports that a temperature upshift within the CHO culture range increases galactosylated glycan species and decreases afucosylated glycan (Marschall et al., 2023). In that study, an upshift from 35.8 °C to 37 °C increased galactosylated species by up to 4.5% and decreased total afucosylated glycan by 1.2%, and temperature is among the few process parameters reported to produce galactosylation shifts exceeding 10% in several cases across the surveyed literature (Bravo-Venegas et al., 2026). However, the present dataset also shows a concurrent, comparably sized rise in Man5, a high-mannose, incompletely processed species, which was not a feature of the more modest 35.8 °C→37 °C upshift (Marschall et al., 2023). This suggests that the 37 °C→39 °C shift applied here, which moves culture temperature above the standard physiological baseline rather than toward it, produced a genuine heat-stress response overlaid on the galactosylation-promoting effect of the upshift itself, rather than a purely beneficial optimization of the glycosylation profile.

The earlier onset and greater magnitude of the G0F decline under glucose+galactose+MnCl feed indicate that the galactose-and manganese-driven substrate effect on the galactosylation pathway, established in the corresponding standard-temperature comparison, is not simply additive with the temperature-upshift effect but appears to accelerate and amplify it. Under standard temperature, the same feed produced a sustained but comparatively gradual separation from glucose feed; here, under concurrent heat stress, the separation in G0F was already substantial within three days of the temperature shift and continued to widen for the remainder of the culture. This amplification also carried a productivity cost that was not observed at standard temperature. In the corresponding standard-temperature comparison at VVD 1.2 (Section 3.4), glucose+galactose+MnCl feed matched or exceeded glucose-feed titer for much of the culture. Under temperature upshift, glucose+galactose+MnCl feed titer remained below glucose feed at every timepoint, with both conditions declining steadily as culture progressed. This suggests that the combination of elevated temperature and galactose+MnCl supplementation imposes a productivity burden that neither factor produces in isolation at these magnitudes, consistent with reports that CHO clones can show nonlinear, condition-dependent productivity responses to temperature shift depending on concurrent process and media conditions (McHugh et al., 2020). Taken together, these results indicate that the galactosylation-enhancing effect of glucose+galactose+MnCl supplementation, previously shown to be productivity-neutral or favorable at VVD 1.2 and standard temperature, does not transfer cleanly to a temperature-upshifted process; both the glycosylation shift and the productivity cost appear to be substantially amplified under concurrent heat stress, an interaction that would need to be accounted for in any process design combining temperature control and galactose-based glycoengineering strategies.

Following the downward temperature shift to 35 °C, the two feeding strategies produced divergent glycoform profiles over the Day 8–20 monitoring window. G0F remained the dominant species under both conditions but followed opposing trajectories: under glucose-19 only feed, G0F rose from approximately 49% on Day 8 to a sustained 57–60% through Day 14, before increasing further to 62–66% by Days 17–19. In contrast, under glucose+galactose+MnCl feeding, G0F declined from a comparable Day 8 value (∼51%) to a plateau of 40–45% from Day 13 onward, resulting in a separation of roughly 20 percentage points between conditions by the end of the culture. This decline in G0F under galactose supplementation was mirrored by a corresponding increase in mono-galactosylated G1F, which rose from ∼12% on Day 8 to a peak of ∼25% on Day 13 and remained elevated (21– 24%) through Day 20. G1F in the glucose-only condition, by comparison, stayed largely flat between 9% and 13% across the same window. A similar but more modest pattern was observed for G1F’, which increased under galactose feeding toward the later culture period (reaching ∼15% by Days 19–20) while declining gradually under glucose-only feeding (from ∼11% to ∼7%). G2F, the digalactosylated species, remained low (<8%) in both conditions, with a marginal upward trend under galactose supplementation that did not reach the magnitude seen for G1F.

**Figure 6.**
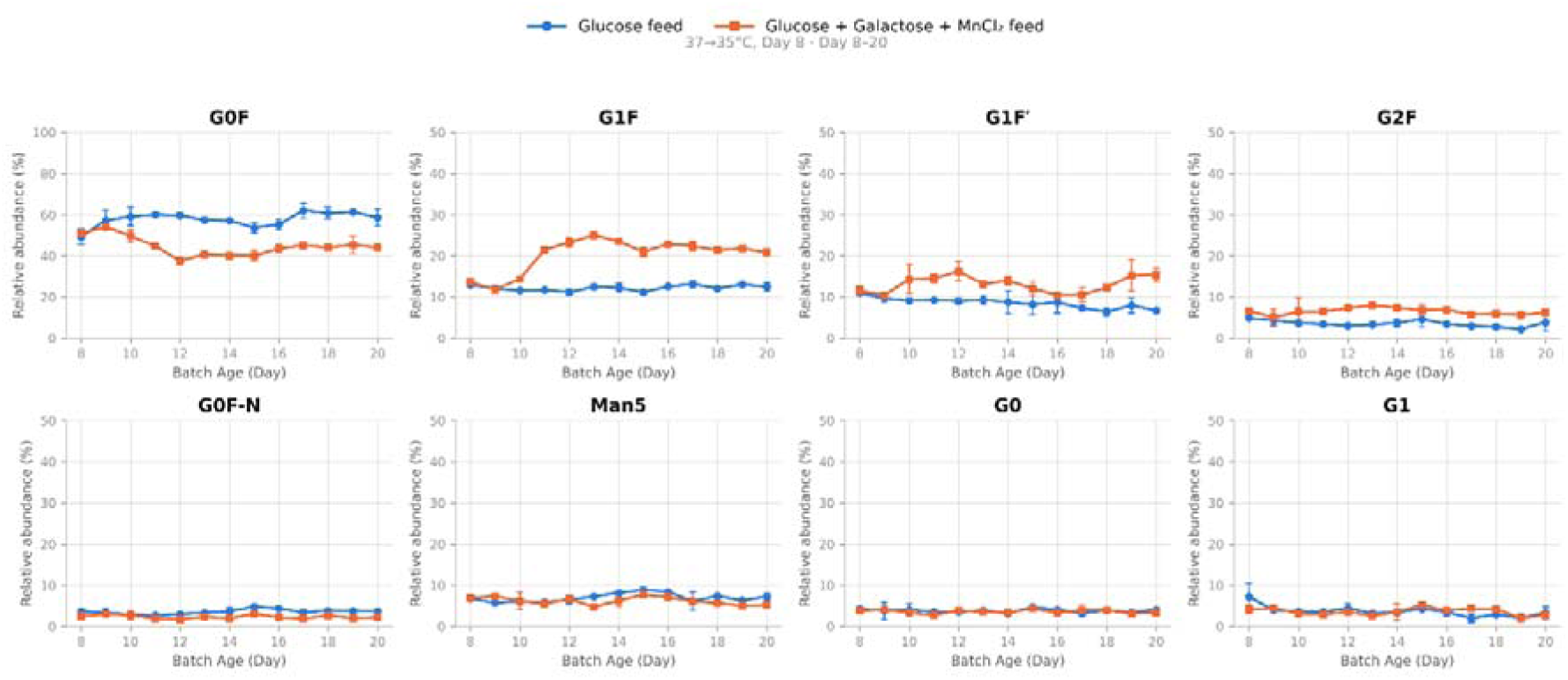
Effect of feed composition on N-glycan relative abundance under temperature downshift. Relative abundance (%) of G0F, G1F, G1F′, G2F, G0F-N, Man5, G0, and G1 over culture duration (Batch Age, Day 8–20) following a temperature shift from 37°C to 35°C on Day 8, comparing glucose feed (blue) versus glucose + galactose + MnCl feed (orange). Points represent mean ± SD of biological duplicates.

Among the minor glycoforms, Man5 was the most abundant species in both conditions (2– 9%) and trended modestly higher under glucose-only feeding after Day 11, peaking at 8.7% on Day 15 versus 7.6% in the glucose+galactose+MnCl condition on the same day. G0F-N, G0, and G1 remained low in relative abundance (generally 2–5%) with substantial overlap between conditions and wide error bars at several timepoints (notably G1 on Day 8), precluding clear separation attributable to feed composition.

Glucose availability was non-limiting under both feed conditions throughout the Day 8–20 window, remaining in a 6.9–14.3 g/L range under glucose feed (n = 2 bioreactors, averaged) and a 7.0–11.0 g/L range under glucose+galactose+MnCl feed (n = 2 bioreactors through Day 12, n = 1 thereafter); neither condition approached the glucose-limited regime seen in the cell density, 1.5 VVD, or 1.2 VVD comparisons. Lactate declined steadily under both conditions over the culture, from 1.4–3.2 g/L (Days 8–9) to 0.6–1.3 g/L (Days 15–20), consistent with the reduced glycolytic flux expected at the lower culture temperature. Ammonia was transiently elevated early in culture under glucose feed, reaching 9.1 mM on Day 10 before declining; from Day 12 onward, ammonia settled into a comparable 1.3–2.9 mM range under both feed conditions, with no sustained feed-dependent separation of the kind observed at 1.5 VVD (Karst et al., 2017). Volumetric titer favored glucose feed through the first half of the monitored window, averaging 0.25–0.38 g/L across Days 9–15 versus 0.18–0.28 g/L under glucose+galactose+MnCl feed. This gap narrowed and then reversed over the second half of culture: glucose-feed titer declined steadily from Day 15 onward, reaching 0.14 g/L by Days 19–20, with one of the two replicates falling substantially faster than the other, while glucose+galactose+MnCl -feed titer stabilized and modestly exceeded glucose feed from Day 17 onward (0.21–0.26 g/L vs. 0.14–0.21 g/L) (Supplementary Fig. 5).

The reciprocal shift from G0F toward G1F and, to a lesser extent, G1F’ and G2F under galactose supplementation is consistent with increased flux through the galactosyltransferase-mediated branch of the N-glycosylation pathway, likely driven by expanded intracellular UDP-galactose availability, in line with prior reports of galactose feeding effects in CHO culture (Gramer et al., 2011). The magnitude of this shift, a roughly 20-percentage-point divergence in G0F by late culture, indicates that galactose supplementation exerted a substantial effect on glycoform distribution even under the reduced-temperature condition, suggesting the two perturbations did not simply offset one another. The comparatively small and inconsistent separation in Man5 and other minor, non-galactosylated species between feed conditions suggests that galactose supplementation primarily influenced later-stage Golgi galactosylation rather than earlier mannose-trimming steps, which may be more sensitive to temperature than to feed composition. The wide error bars observed for several minor glycoform timepoints (e.g., Day 8 G1, Day 12 G0F) reflect biological replicate variability rather than a systematic feed or temperature effect and should be interpreted with that caveat when comparing conditions at individual timepoints.

The non-limiting glucose availability sustained under both feed conditions throughout this comparison isolates the G0F-to-G1F/G1F’ shift as a feed-composition effect rather than a secondary consequence of glucose restriction, a confound that could not be excluded in the cell density, 1.5 VVD, or 1.2 VVD comparisons. The comparable ammonia trajectories between conditions from Day 12 onward further argue against an ammonia-mediated contribution of the kind proposed for the 1.5 VVD comparison (Karst et al., 2017), reinforcing a direct substrate-driven mechanism as the primary driver of the glycoform separation observed here. The late-culture titer crossover, with glucose-feed titer declining from Day 15 onward while glucose+galactose+MnCl -feed titer stabilizes, is a more tentative finding than the glycoform results. It is driven in large part by a steep decline specific to one glucose-feed replicate rather than a trend shared across both, and the glucose+galactose+MnCl -feed condition is represented by a single reactor from Day 13 onward. The apparent productivity advantage for the combined feed in late culture should therefore be treated as an observation to confirm with additional replicates rather than a robust conclusion, in contrast to the better-replicated titer comparisons at VVD 1.2 and VVD 1.5.

### 3.6 Impact of pH on Trastuzumab N-glycosylation

The two pH shift conditions produced markedly divergent glycoform profiles, with the pH shift down condition (6.7) showing a pronounced departure from baseline, with titer and glycan data no longer quantifiable from Day 17 onward. Under pH shift up (7.5), G0F remained the dominant species and stayed relatively stable across the full 20-day window, ranging between 49% and 59% with a modest decline through the mid-culture period followed by a return to ∼51–52% by Day 20. Under pH shift down, G0F instead rose sharply from a comparable Day 8–11 baseline (∼57%) to a peak of ∼76% on Day 13, remained elevated through Day 14 (∼75%), and then declined to ∼62% by Day 16, the last timepoint at which glycan data could be generated for this condition.

This transient G0F increase under pH shift down coincided with a corresponding depletion of galactosylated species. G1F declined steeply from ∼20% on Day 8 to ∼6% by Day 12, and G1F’ followed a similar trajectory, falling from ∼12% to ∼6% over the same interval; both remained suppressed through the final measured timepoints. In contrast, G1F and G1F’ under pH shift up declined only gradually across the full culture, settling at ∼15% and ∼11%, respectively, by Day 20. G2F remained low (<7%) and broadly comparable between conditions throughout. Among the minor glycoforms, the pH shift down condition showed the most substantial deviation in Man5, which increased steadily from ∼4% on Day 8 to ∼24% by Day 16, the largest relative change observed across any glycoform in either condition. G1 followed a comparable late-culture rise under pH shift down, reaching ∼14% by Day 16 with wide variability (error bars spanning approximately 6–20% at that timepoint). Under pH shift up, both Man5 and G1 remained low and stable throughout (generally 3–7%), tracking closely with G0 and G0F-N, which showed similarly low trajectories in both conditions.

**Figure 7.**
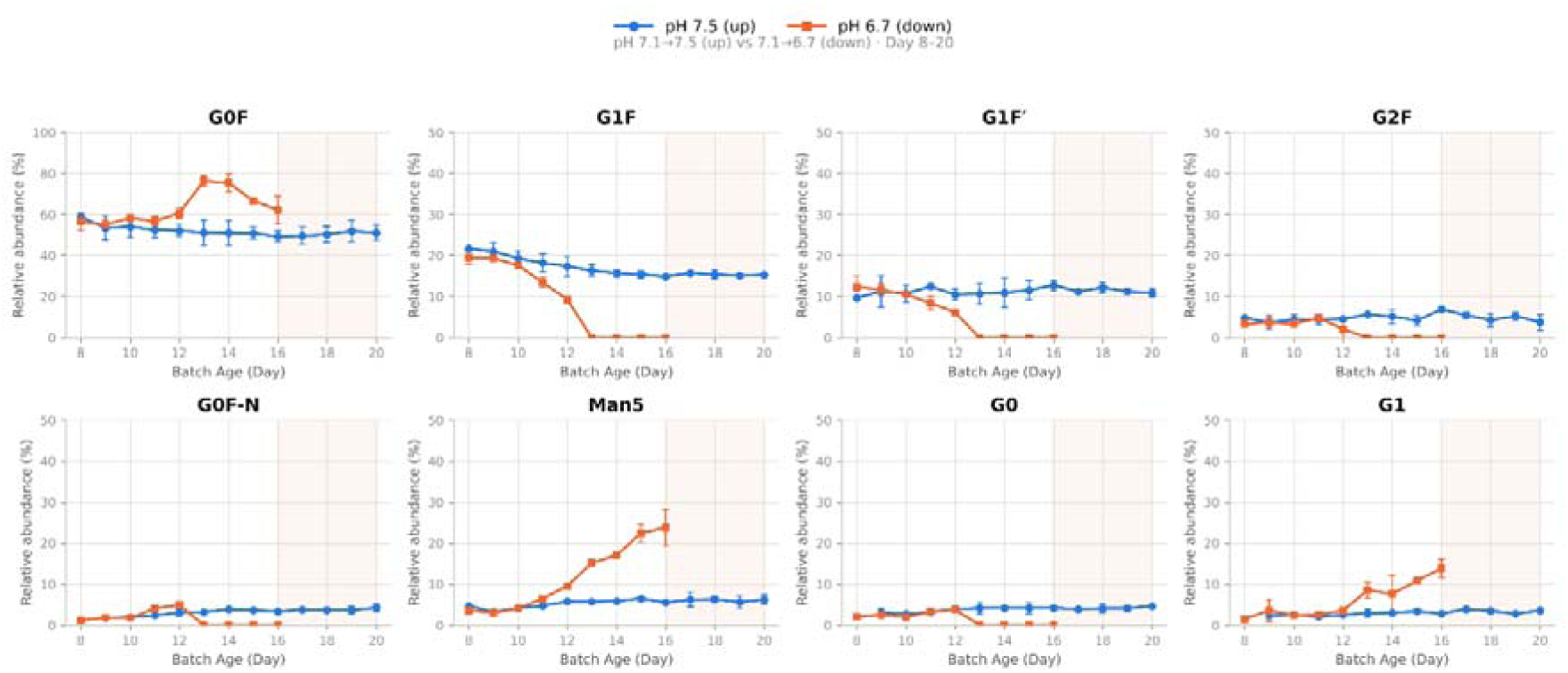
Effect of pH shift direction on N-glycan relative abundance. Relative abundance (%) of G0F, G1F, G1F′, G2F, G0F-N, Man5, G0, and G1 over culture duration (Batch Age, Day 8–20), comparing pH upshift to 7.5 (blue) versus pH downshift to 6.7 (orange). Data for the pH-downshift condition are absent beyond Day 16–17 due to culture termination. G0F is plotted on a 0–100% axis; all other glycoforms are plotted on a 0–50% axis. Points represent mean ± SD of biological duplicates.

Glucose availability diverged sharply between the two pH conditions. Under pH shift up, glucose was actively consumed throughout, declining from 6.4–6.7 g/L to a glucose-limited range (<1 g/L) by Days 15–16 and reaching near-depletion (0.1–0.2 g/L) by Days 17–20. Lactate rose correspondingly, from 1.9–2.3 g/L to 3.4–5.4 g/L through the mid-to-late culture period, consistent with sustained glycolytic activity. Under pH shift down, glucose uptake effectively ceased from approximately Day 10 onward: rather than declining, glucose concentration rose steadily across the remainder of the culture, from 6.2–6.6 g/L (Days 8–9) to 12.3–13.3 g/L by Day 20, as feed continued to be supplied without corresponding consumption. Lactate collapsed in parallel, falling from 2.4–2.5 g/L to below 0.3 g/L from Day 11 onward and remaining at this floor through Day 20. Viable cell density diverged markedly between conditions from Day 11 onward. Under pH shift up, VCD remained high and stable throughout, ranging from 16.1–28.7 × 10 cells/mL with viability consistently above 99%. Under pH shift down, VCD declined steadily from a comparable peak (24.2–31.9 × 10 cells/mL, Days 9–11) to 6.8–8.5 × 10 cells/mL by Days 15–16, and fell further to 1.1–4.8 × 10 cells/mL by Days 17–20. Viability declined only modestly over this window (92.5–97.6%), indicating a sustained decline in cell density rather than an acute viability collapse. Ammonia was modest and comparatively stable under pH shift up (generally 0.6– 1.5 mM, with isolated spikes to 2.1–4.1 mM on Day 19). Under pH shift down, ammonia rose transiently early in the shift, peaking at 3.6–4.3 mM over Days 11–13, before declining steadily to 0.9–1.6 mM by Days 17–20 as cell density fell. Volumetric titer tracked the divergent VCD trajectories of the two conditions closely. Under pH shift up, titer rose over the culture period, from 0.19–0.21 g/L (Days 8–9) to a peak of 0.50 g/L, tracking the sustained high cell density. Under pH shift down, titer declined steadily from a comparable starting point (0.14–0.25 g/L, Days 8–10) to 0.023–0.026 g/L by Day 16 as VCD fell below 10 × 10 cells/mL; from Day 17 onward, as VCD fell further to 1–5 × 10 cells/mL, the UPLC elution peak was no longer integrable in either reactor, consistent with a cell population too sparse to generate a detectable secreted-product signal rather than a loss of per-cell productivity (Supplementary Fig. 6).

The pronounced rise in G0F and Man5 alongside depletion of G1F and G1F’ under acidic pH shift (6.7) is consistent with impaired galactosyltransferase activity and disrupted glycan trimming under sub-physiological Golgi and ER pH, both of which are known to be sensitive to extracellular and intracellular pH shifts in CHO culture (Ivarsson et al., 2014). The concurrent rise in Man5, an under-processed high-mannose intermediate, suggests that the low-pH condition affected glycan maturation upstream of galactosylation as well, rather than acting solely at the level of terminal galactose addition. This pattern differs from the galactose-feed effect observed in the temperature downshift comparison, where G1F increased at the expense of G0F without a comparable rise in Man5, pointing to a distinct mechanism, likely enzymatic disruption from pH stress rather than substrate-driven pathway flux. The decline in VCD under pH shift down, from a peak comparable to that under pH shift up to 1–5 × 10 cells/mL by Days 17–20, indicates a sustained growth and survival defect under acidic stress rather than an acute culture failure: viability remained relatively high throughout even as VCD fell, suggesting cells were being lost gradually rather than undergoing a rapid viability collapse. This VCD decline likely explains the loss of integrable titer from Day 17 onward: at cell densities below approximately 5 × 10 cells/mL, secreted product in the perfusate likely fell below the assay’s detection threshold, independent of any change in per-cell productivity. Glucose, lactate, and ammonia continued to be measured through Day 20 under this condition because these are perfusate-level measurements that do not require a minimum cell density to register, confirming that the bioreactor continued to operate throughout the monitored window even as the cell population, and with it the quantifiable titer and glycan signal, diminished. This asymmetry between the two pH conditions is itself a relevant process-control finding which suggests that upward pH excursions are comparatively well tolerated, whereas downward excursions of a similar magnitude impose a sustained growth defect that compromises both product quantifiability and glycosylation-based quality assessment well before culture operation itself is affected.

### 3.7 Cross-condition comparison: a decision logic for process-parameter selection and process perturbation recovery strategy

The preceding sections (3.2–3.6) each addressed a single process lever in isolation, such as cell density, feed composition, perfusion intensity, temperature, and pH, with each evaluated against its own glycosylation, productivity, and cell-health outcomes. Taken together, these six comparisons do not point to a single “best”1 condition but to a set of trade-offs that a process-design decision has to weigh explicitly. Figure 8 consolidates this experimental evidence into a single decision-logic diagram, tracing each perturbation through its observed glycosylation shift, titer impact, and cell-health response to a qualitative risk tier (favorable, moderate/promising, trade-off, or avoid/severe), without invoking any computational or mechanistic model Each classification is derived directly from the glycoform, metabolite, titer, and VCD data presented in Sections 3.2–3.6, in contrast to model-supported approaches to glycosylation control in perfusion culture, in which glycoform trajectories are predicted from mathematical or flux-based models (Zhang et al., 2020).

**Figure 8.**
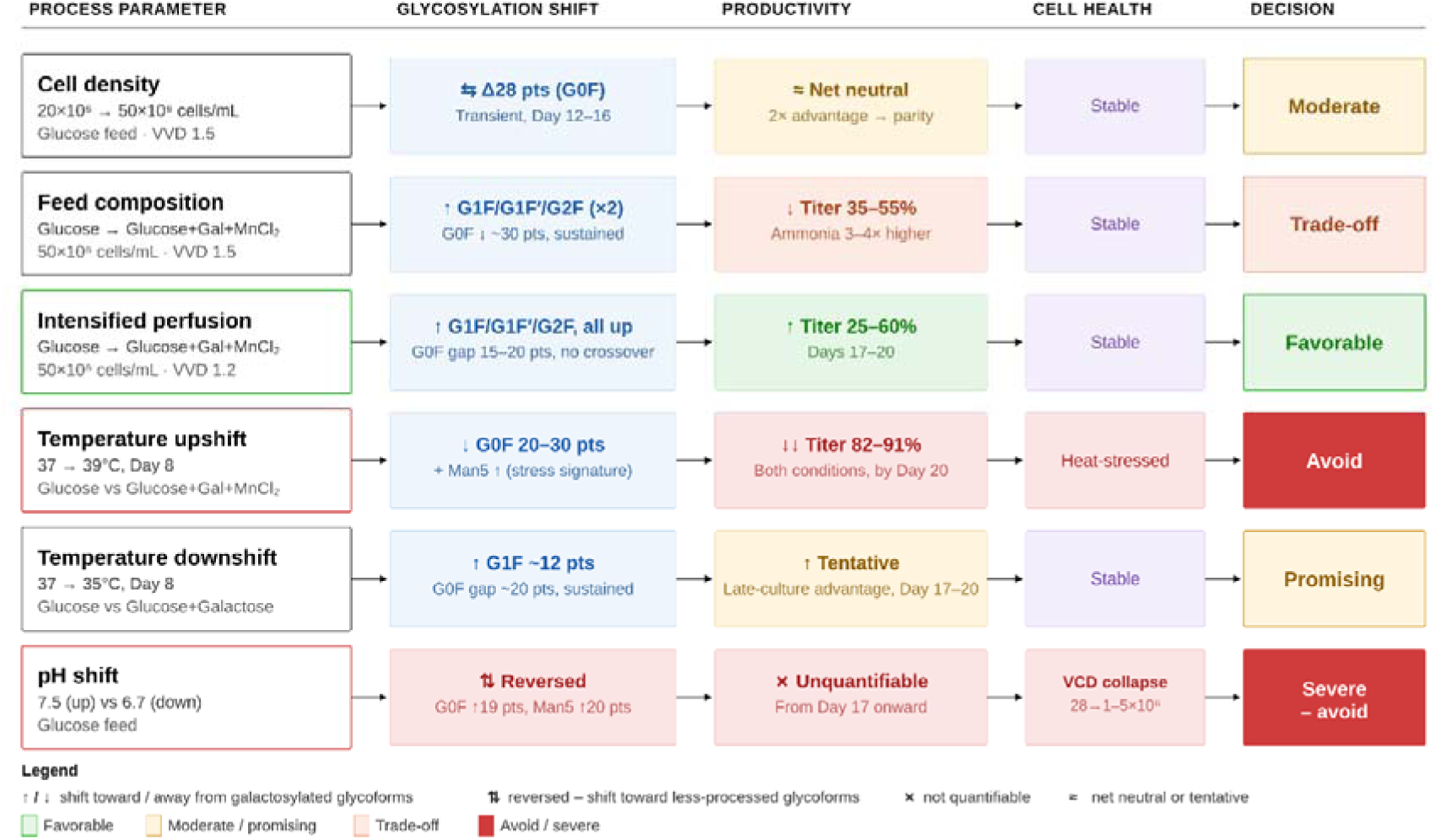
Decision logic for six process-parameter comparisons (Sections 3.2–3.6) in Trastuzumab-producing CHO microperfusion culture. Each row traces the perturbation through its observed glycosylation shift, productivity (titer) impact, and cell-health impact to a process decision, built directly from the reported experimental glycoform, metabolite, titer, and VCD data; no computational or mechanistic modeling. Box shading in the Decision column indicates risk tier from favorable (open) to avoid/severe (filled).

Two conditions clearly separate from the rest. Intensified perfusion with the combined glucose–galactose–MnCl feed at 1.2 VVD reproduced the desired, sustained shift toward galactosylated glycoforms (G1F, G1F′, G2F) seen under the combined feed at 1.5 VVD, but without the earlier titer penalty, with productivity matching or exceeding the glucose-only control by Day 17–20, and cell health was not compromised. This condition is therefore the only one in the panel that achieves the glycoengineering objective at no measurable productivity cost, and it is flagged as the favorable outcome of the study. At the opposite end, temperature upshift to 39°C and pH shift to 6.7 were both associated with severe, compounding failure. The former combined an 82–91% titer collapse with a Man5-co-elevated glycosylation signature consistent with heat stress rather than a controlled glycoengineering effect; the latter combined an unquantifiable late-culture titer with a near-total collapse in viable cell density (≈28 → 1–5 × 10 cells/mL) and a reversal of the glycosylation shift toward less-processed forms. Both are classified as conditions to avoid under the tested regime.

The remaining four comparisons occupy intermediate ground and illustrate that “favorable” and “effective” are not interchangeable. Feed composition alone (at 1.5 VVD, without the perfusion-rate reduction) achieved the same directionally desired glycosylation shift as the intensified-perfusion condition, but at a consistent 35–55% titer cost and markedly elevated ammonia, a trade-off acceptable only if quality attributes are prioritized over volumetric productivity. Temperature downshift to 35°C produced a comparable glycosylation benefit with a tentative, late-emerging productivity advantage. Cell density alone, independent of feed or perfusion rate, produced only a transient glycosylation crossover that reconverged by Day 19–20 and a titer advantage that eroded to parity, consistent with reports that cell density itself had little influence on N-glycosylation at comparable cell-specific perfusion rate(Schwarz et al., 2022), so this condition is best read as process-neutral rather than as a lever for glycoform control.

The temperature- and pH-shift results also carry a practical implication for how an in-process excursion should be managed, and the two stressors are not equally recoverable. pH downshift is the only condition in this panel associated with a collapse of the viable cell population itself (≈28 → 1–5 × 10 cells/mL by Day 17–20), rather than a productivity or glycosylation defect superimposed on an intact population. A culture reduced to this viable-cell fraction is unlikely to recover regardless of when pH is subsequently corrected, so pH downshift is best treated as a hard, non-recoverable failure mode: the appropriate control action is a predefined abort/discard criterion (e.g., sustained pH below the downshift setpoint, or VCD falling below a defined threshold) rather than a reversion-and-rescue protocol, consistent with the use of predefined control actions and design-space boundaries to keep a process in a state of control (2004).

Temperature upshift, although equally severe in its titer and glycosylation-quality impact, was characterized in this dataset as a heat-stressed rather than a collapsed population, implying a metabolic and productivity impact superimposed on cells that remained viable. This distinction is the basis for treating it, and by the same logic pH upshift, as potentially recoverable if the intervention is timed correctly. Because titer and glycoform changes lag behind the underlying stress response, a rescue strategy should trigger on earlier indicators of stress onset such as rising lactate/ammonia, declining specific productivity, or an increase in the Man5 stress signature rather than waiting for titer or VCD to visibly turn over, by which point the Day 12–16 transient-crossover behavior observed under the cell-density comparison suggests the window for a clean reversion has likely closed. A graded return to the 37°C setpoint, paired with a short-term feed boost (glucose, or the combined glucose+galactose+MnCl feed already shown in this study to support titer under comparable conditions), is hypothesized to be more defensible than an abrupt step change or reversion alone. Temperature downshift, by contrast, did not compromise cell health or productivity in this dataset and so does not require a rescue protocol; where downshift needs to be reverted, it is a scheduling decision rather than a recovery intervention. For pH upshift, the same early-trigger logic as temperature upshift is proposed as a working assumption in the absence of population-level data specific to that arm, and should be flagged as such rather than treated as an established result.

Read together, these findings indicate that durable, desired shifts in N-glycosylation in this microperfusion system are feed- and perfusion-rate–dependent rather than density- or temperature-dependent, that the 1.2 VVD perfusion-rate reduction appears to be the modification that converts a glycosylation-beneficial feed strategy from a productivity trade-off into a net-favorable one, and that thermal and pH excursions may require fundamentally different control responses, including early biomarker-triggered rescue for stress-type failures and a hard discard criterion for population-collapse failures such as pH downshift. This decision logic and its associated control strategy are intended as a practical reference for prioritizing which of the tested interventions merit adoption, further replicate confirmation, or exclusion, and for setting in-process intervention criteria, in subsequent microperfusion process development.

## Conclusion

This study applied the N-GLYcanyzer PAT workflow to resolve time-resolved N-glycosylation dynamics in Trastuzumab-producing CHO microperfusion cultures across five systematically varied process parameters: carbon source, cell density, perfusion rate, temperature, and pH. Across these comparisons, nucleotide-sugar donor and cofactor availability, principally glucose status and galactose/MnCl supplementation, consistently governed the balance between G0F and the more extensively processed glycoforms (G1F, G1F’, G2F), while temperature and pH perturbations acted through largely independent pathways that compounded, compounding high-mannose accumulation and productivity loss rather than substituting for the substrate-driven effect. The galactose/MnCl feed strategy achieved a sustained, productivity-neutral galactosylation shift under standard temperature and pH, but this effect was amplified in both magnitude and productivity cost under concurrent heat stress, and an acidic pH shift produced the most severe outcome observed, a progressive decline in viable cell density that rendered titer and glycan data unquantifiable well before culture metabolism itself was disrupted.

These findings carry direct relevance to process control strategy as continuous and intensified biomanufacturing platforms move toward attribute-based, rather than purely time- or volume-based, operating regimes (Coffman et al., 2021). Microperfusion systems compress the residence time available to detect and correct a glycosylation excursion relative to fed-batch, and the non-additive interactions observed here among feed composition, temperature, and pH indicate that a design space characterized along individual process parameters, as is common practice when adapting a fed-batch control strategy to a perfusion platform, may underestimate the risk of a CQA excursion at the intersection of two or more parameters (2004). The real-time glycoform resolution demonstrated by the N-GLYcanyzer workflow across all six comparisons supports its use not only for retrospective process characterization, as applied here, but as the measurement layer for a PAT-enabled, attribute-based control loop capable of triggering a feed or setpoint adjustment within a perfusion culture’s operating timescale, consistent with the process analytical technology framework’s original intent of building quality into the process rather than testing for it at the end (Chemmalil et al., 2020). Extending this PAT-monitored framework to higher biological replication, additional product molecules, and pilot- or commercial-scale perfusion systems would be a necessary next step toward establishing it as a validated, scale-independent component of a real-time release and active control strategy for continuous biomanufacturing (Coffman et al., 2021).

Such a framework would also need to define acceptance limits for each glycoform attribute and document how the PAT measurements map onto those limits across batches, so that regulatory expectations for demonstrated process understanding can be met in a perpetual, non-batch operating mode (Reusch and Tejada, 2015). Furthermore, incorporating these real-time analytics into a feedback loop necessitates the validation of automated data-handling protocols to ensure that high-frequency glycan resolution is robust against sensor drift or signal noise inherent to long-term, steady-state cultivation.

## Acknowledgements

This work was supported by awards #PC5.2-112 and #PC7.1-146 granted to Rutgers University under a Project Award Agreement from the National Institute for Innovation in Manufacturing Biopharmaceuticals (NIIMBL) and financial assistance award 70NANB21H086 from the US Department of Commerce, National Institute of Standards and Technology. During the preparation of this manuscript, the authors used Claude (Anthropic) to improve the layout, typography and visual clarity of Figure 1 and Figure 8, which were originally created using BioRender. The authors also used Gemini (Google) for assistance with grammatical checking and sentence flow refinement. Following the use of these tools, the authors reviewed, edited, and verified all content to ensure accuracy, scientific integrity, and fidelity to the original data, and take full responsibility for the final published content.

## Declaration of competing interest

Rutgers/SPSC has a pending patent filed for the N-GLYcanyzer (US20240110925A1). The authors declare that they have no known competing financial interests or personal relationships that could have appeared to influence the work reported in this paper.

