## Supplementary Information (SI) for "N-GLYcanyzer monitoring of a microperfusion process reveals critical parameters’ impact on Trastuzumab biosimilar N-glycoform time evolution"

### Supplementary figures

#### 1) Metabolite and titer profiles accompanying the N-glycosylation comparisons in Figures 2–7.

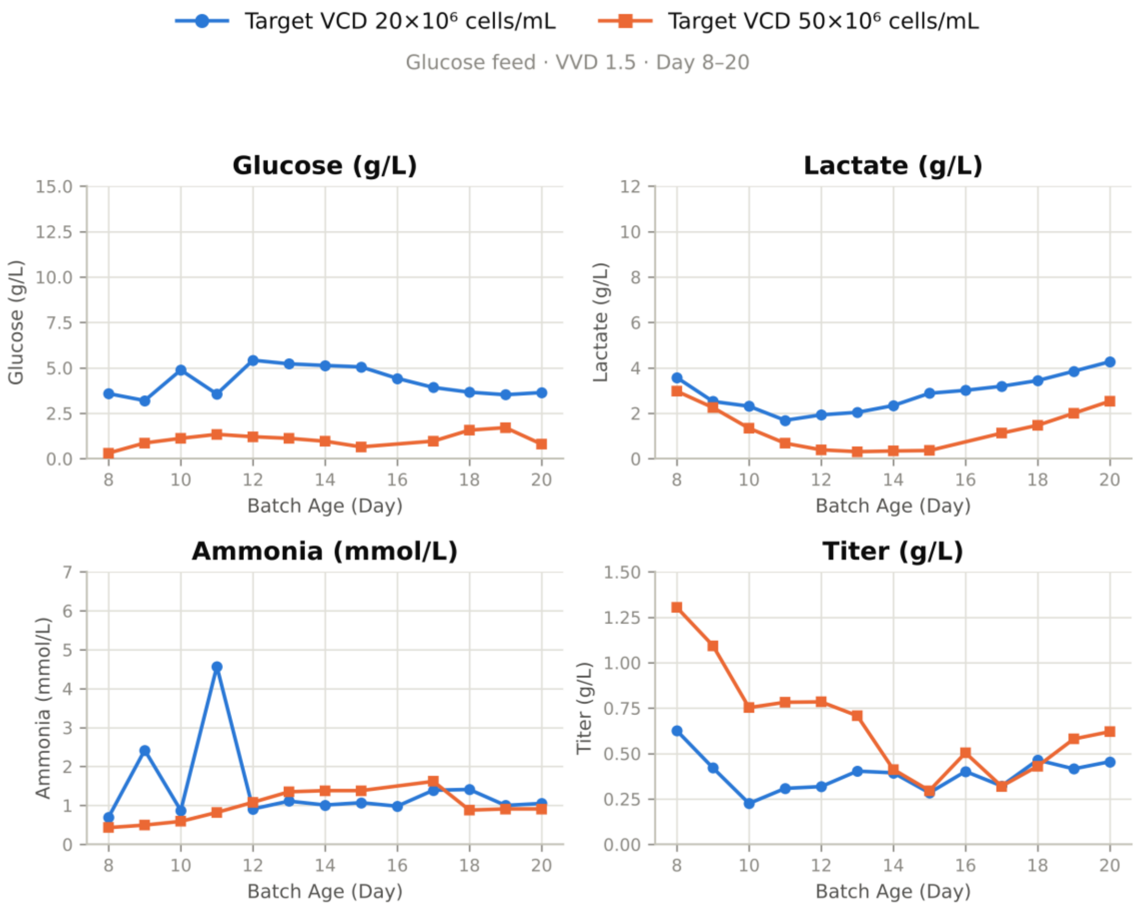


**Figure S1.** Metabolite and titer profiles for the cell-density comparison (companion to Figure 2). Glucose-fed cultures at 1.5 VVD were controlled to a target viable cell density of 20×10⁶ cells/mL (blue) or 50×10⁶ cells/mL (orange). Panels show glucose (g/L), lactate (g/L), ammonia (mmol/L) and volumetric titer (g/L) over culture duration (Batch Age, Day 8–20).

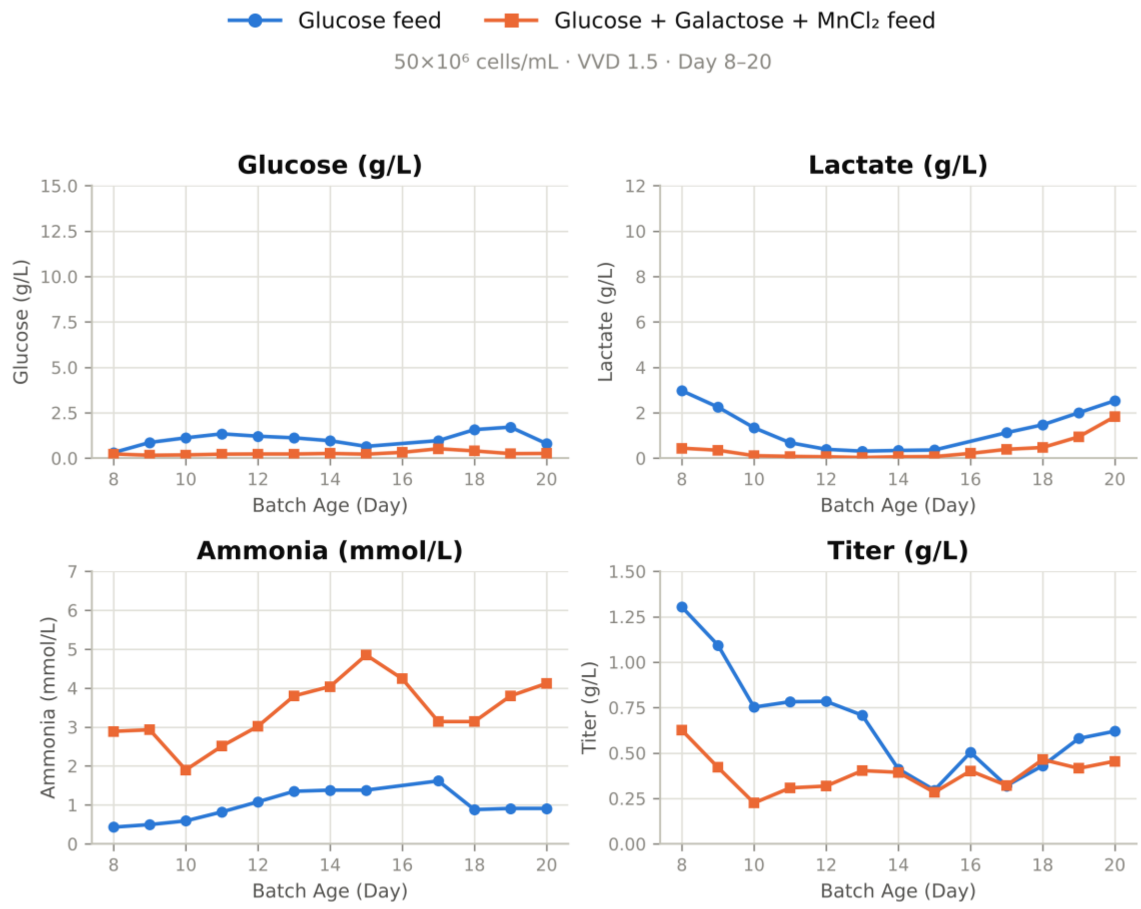


**Figure S2.** Metabolite and titer profiles for the feed-composition comparison at controlled cell density (companion to Figure 3). Cultures at 50×10⁶ cells/mL and 1.5 VVD received glucose feed (blue) or glucose + galactose + $MnCl_2$ feed (orange). Panels show glucose (g/L), lactate (g/L), ammonia (mmol/L) and volumetric titer (g/L) over culture duration (Batch Age, Day 8–20).

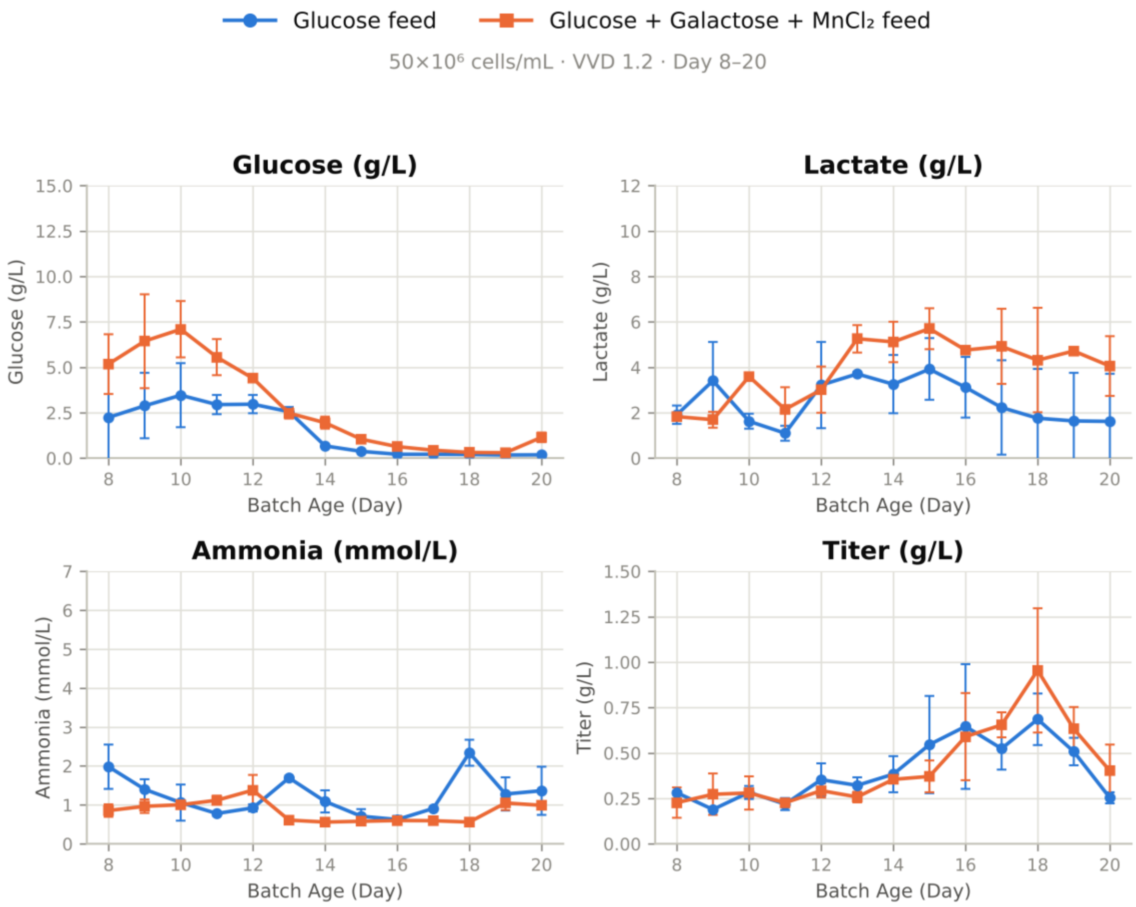


**Figure S3.** Metabolite and titer profiles for the feed-composition comparison under intensified perfusion (companion to Figure 4). Cultures at 50×10⁶ cells/mL and 1.2 VVD received glucose feed (blue) or glucose + galactose + $MnCl_2$ feed (orange). Panels show glucose (g/L), lactate (g/L), ammonia (mmol/L) and volumetric titer (g/L) over culture duration (Batch Age, Day 8–20).

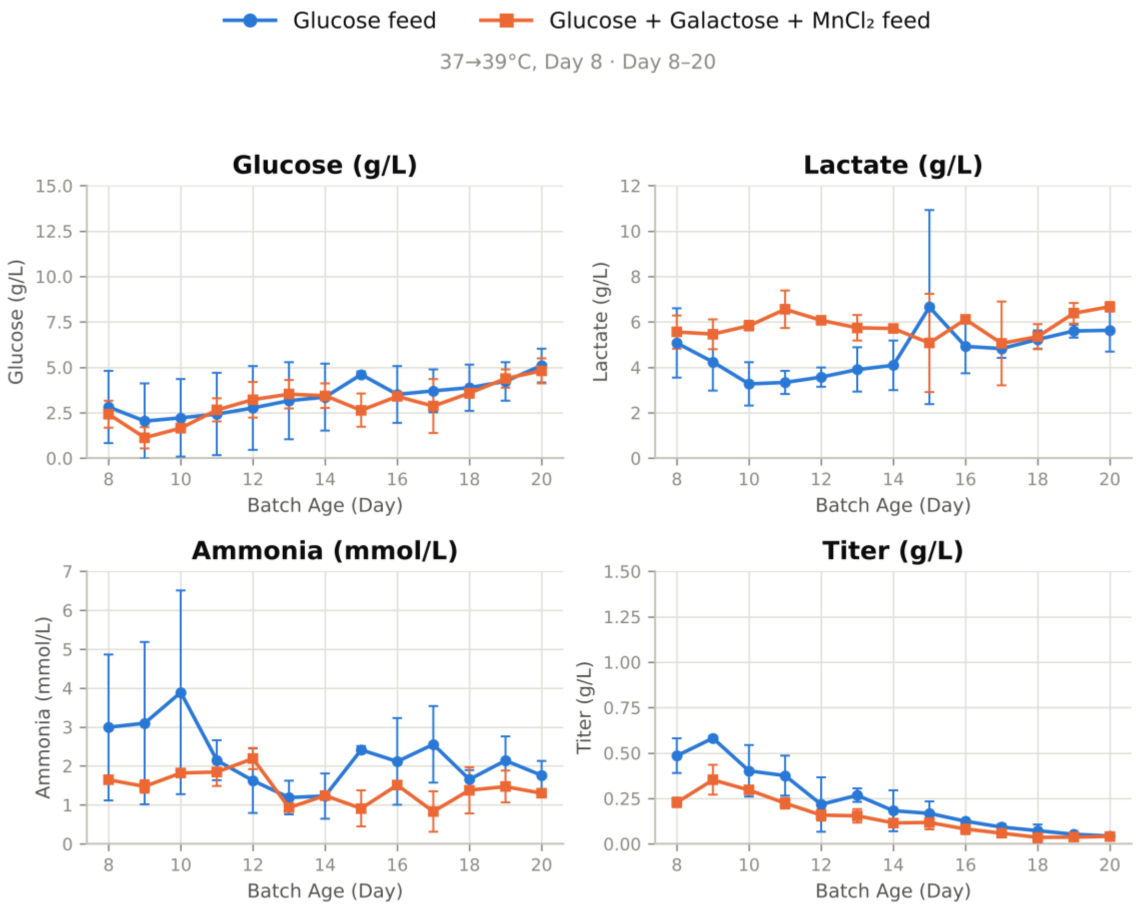


**Figure S4.** Metabolite and titer profiles for the feed-composition comparison under temperature upshift (companion to Figure 5). Cultures were shifted from 37 to 39 °C on Day 8 and received glucose feed (blue) or glucose + galactose + $MnCl_2$ feed (orange). Panels show glucose (g/L), lactate (g/L), ammonia (mmol/L) and volumetric titer (g/L) over culture duration (Batch Age, Day 8–20).

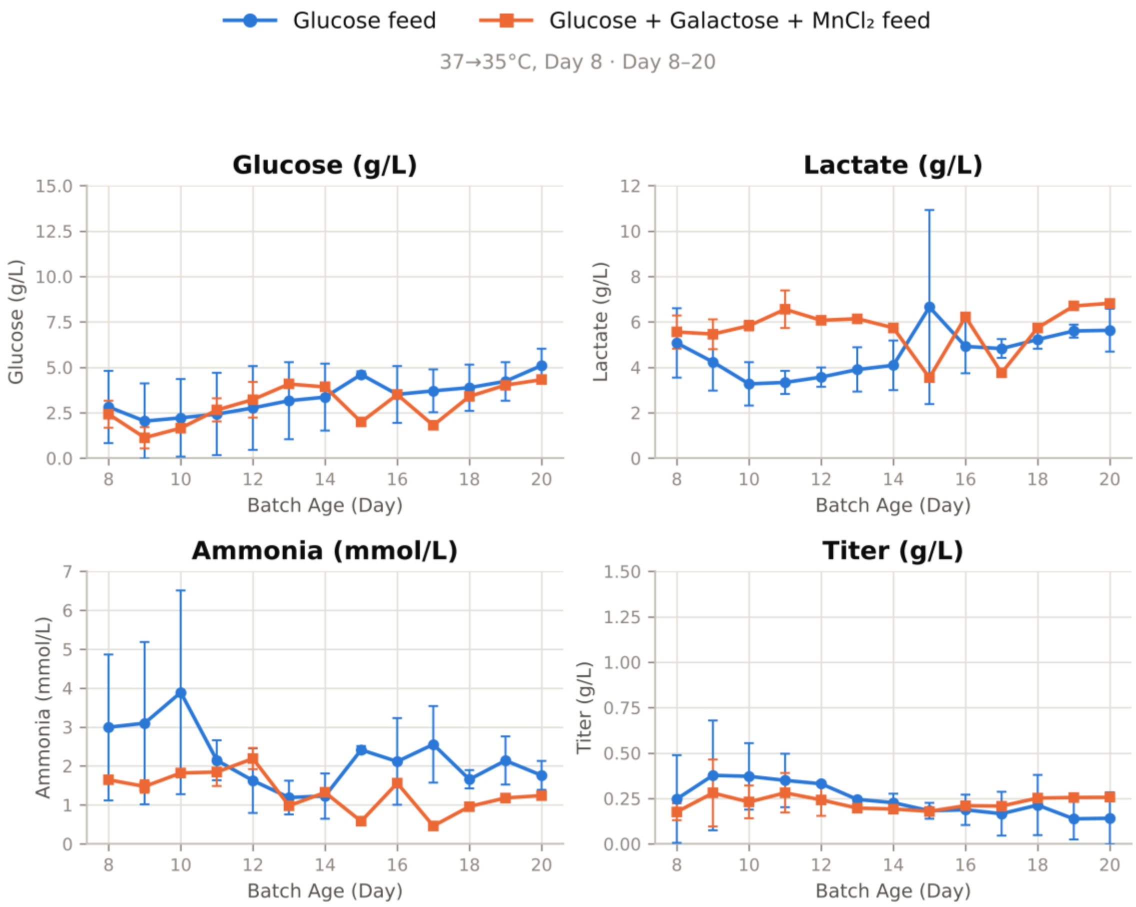


**Figure S5.** Metabolite and titer profiles for the feed-composition comparison under temperature downshift (companion to Figure 6). Cultures were shifted from 37 to 35 °C on Day 8 and received glucose feed (blue; mean of two bioreactors) or glucose + galactose + $MnCl_2$ feed (orange). Panels show glucose (g/L), lactate (g/L), ammonia (mmol/L) and volumetric titer (g/L) over culture duration (Batch Age, Day 8–20).

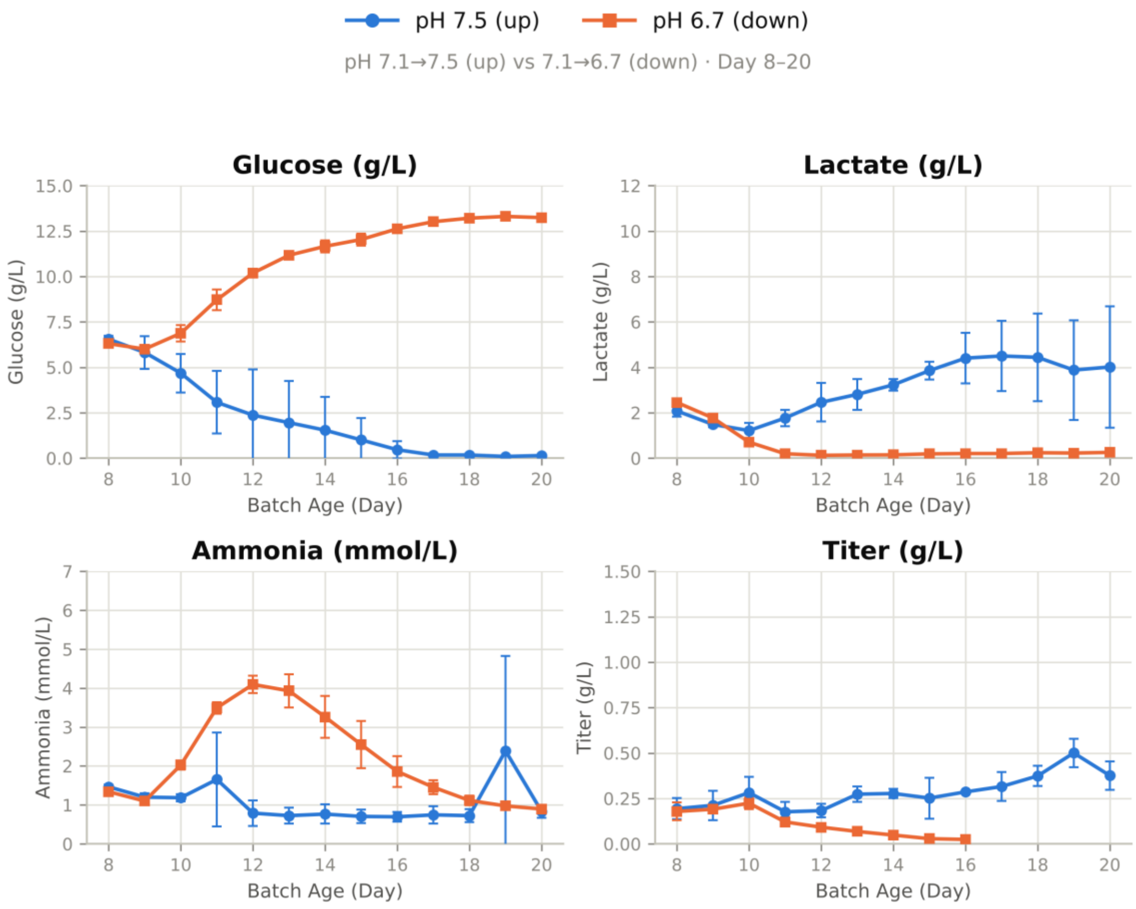


**Figure S6.** Metabolite and titer profiles for the pH-shift comparison (companion to Figure 7). Cultures were shifted from pH 7.1 to 7.5 (blue) or to 6.7 (orange). Titer for the pH 6.7 condition is not reported from Day 17 onward, consistent with the loss of culture viability in this condition. Panels show glucose (g/L), lactate (g/L), ammonia (mmol/L) and volumetric titer (g/L) over culture duration (Batch Age, Day 8–20).
